# High-Throughput Fragment Screening Identifies a New Small Molecule Scaffold that Modulates TREM2 Signaling

**DOI:** 10.1101/2025.11.23.690065

**Authors:** Natalie Fuchs, Farida El Gaamouch, Hossam Nada, Sungwoo Cho, Moustafa T. Gabr

## Abstract

Fragment-based drug discovery (FBDD) remains a powerful tool in drug development for targeting a wide range of proteins and identifying new small molecule-based scaffolds. Here, we explored a fragment library of 3,200 compounds using a temperature-related intensity change (TRIC)-based high-throughput screening (HTS) approach, and successfully identified new scaffolds that bind to triggering receptor expressed on myeloid cells 2 (TREM2), a relevant target in neurodegenerative diseases and cancer immunotherapy. We first validated the hits with dose-dependent assays, then chose the three most promising compounds (**2M06**, **6B10**, **7G19**) with binding affinities in the low to medium micromolar range for a “SAR by catalog” study. We screened 29 selected derivatives and subsequently evaluated them with dose-dependent experiments, a thermal shift assay (TSA), selectivity studies with off-targets (LAG-3, TREM1), and finally, in vitro TREM2 activation assays. In this SAR study, derivative **6B10-9** emerged as the lead compound with moderate TREM2 binding affinity (*K*_D_ = 68.3 µM) and significant effects on TREM2-dependent phosphorylation of SYK and DAP12 in HEK cells as wells as on microglial phagocytosis in HMC3 cells. Additionally, an in silico analysis revealed that **6B10-9** forms a stable complex with TREM2 via hydrogen bonding, which maintains its structural integrity during extended molecular dynamics (MD) simulations. These results suggest that **6B10-9** could serve as a promising lead for future optimizations in the development of small molecule-based TREM2 modulators.

## Introduction

Targeting triggering receptor expressed on myeloid cells 2 (TREM2) is not only relevant in neurodegenerative diseases like Alzheimer’s Disease (AD), but also in various tumors (e.g., ovarian cancer, lung cancer).^1^ The immunomodulatory receptor is commonly found on the cell surface of myeloid cells such as microglia or dendritic cells and plays a role in regulating the immune response.^2^ In the central nervous system (CNS), TREM2 is crucial for microglial activation and survival, and is therefore closely associated with neurodegeneration.^3, 4^ In Alzheimer’s Disease, TREM2 supports the microglial amyloid-beta (Aβ) plaque clearance and, on top of that, certain variants (e.g., R47H mutation) have been shown to be associated with a higher risk for disease progression.^5–7^ In peripheral tissues, TREM2 is mainly expressed on dendritic cells and has been found to be overexpressed on tumor-associated macrophages (TAMs).^8^ Here, it contributes to an immunosuppressive tumor microenvironment (TME) through reduced CD8^+^ T cell infiltration and tumor cytotoxicity.^9^ Inhibiting TREM2 leads to reprogramming of TAMs to their pro-inflammatory state and thus to an increased anti-tumor response.^10^ In addition, studies have investigated the potential of combination therapies with anti-TREM2 treatment and PD-1 blockade to combat anti-PD-1-resistant tumors.^11^ However, TREM2 has a protective function in CNS tumor models, highlighting the need for tissue-specific TREM2 ligands as potential cancer immunotherapies or therapeutics in neurodegenerative diseases.^12^ So far, many approaches to target TREM2 have focused on monoclonal antibodies (mAbs) rather than small molecules.^13, 14^ Notable small molecule-based ligands include **VG-3927**,^15^ a TREM2 agonist currently in clinical trials, and Sulfavant A,^16^ the first synthetic molecule to bind TREM2. Since TREM2 is found in multiple tissues and has different functions in each, finding suitable ligands and tailoring them towards specific use in these different diseases is key.

In target-based drug discovery (TBDD), various approaches can help to identify new scaffolds. Commonly used strategies include virtual screenings (VS), the use of DNA-encoded libraries (DEL) or high-throughput screenings (HTS). Fragment-based drug discovery (FBDD) is a powerful and versatile tool to find new scaffolds with multiple advantages over other HTS methods.^17^ For once, the fragments, which are low molecular weight compounds (typically 150–300 Da), have a greater potential for optimization that is still within the scope of traditional “rule-of-five” drug candidates.^18^ However, they usually display weaker binding affinities than other small molecules, mainly because their scaffolds only allow limited interactions in binding sites.^19^ Therefore, a suitable HTS method should be able to detect weak binders (*K*_D_ in medium to high micromolar range) as well.^20^ We recently established an HTS platform for TREM2 based on temperature-related intensity change (TRIC, also known as microscale thermophoresis, MST) that was able to robustly identify weak binders with affinities in the medium to high micromolar range in a proof-of-concept screening.^21^

Here, we applied our screening platform to a fragment library with 3,200 compounds in search of small molecules with new scaffolds to bind TREM2. We followed up with a “SAR by catalog” study for the most promising hits and evaluated them with binding affinity and selectivity studies as well as functional cell assays. Finally, we concluded with an in silico binding analysis for the top compound.

## Materials and Methods

### Compounds

The compounds were purchased in form of a library from Enamine LLC (Monmouth Junction, NJ, USA), namely the “Enamine PPI Fragment Library” (3,200 compounds, #PPIF-3200-Y-10). The library was stored at –80 °C on 384 well plates as 10 mM DMSO stock solutions. Additional derivatives were purchased from Enamine LLC as solids and reconstituted to 50 mM stocks with DMSO. All stocks were aliquoted and stored at –80 °C until use.

### High-Throughput Fragment Screening with Dianthus

#### Initial Single-Dose Screening

The screening was performed using a temperature-related intensity change (TRIC)-based method on Dianthus NT.23Pico (NanoTemper Technologies, Germany) as described previously, with a few adjustments.^21^ His-tagged TREM2 (SinoBiological, #11084-H08H) was labeled with RED-tris-NTA 2^nd^ generation dye (NanoTemper, #MO-L018) according to the manufacturer’s protocol. The libraries were transferred from 384-well DMSO stock plates (10 mM) to intermediate plates as 2-fold ligand stocks (200 µM, PBS pH 7.4, 0.05% Tween-20, 4% DMSO) with the help of an Integra mini-96 pipetting robot. For the screening in a 384-well plate format, 10 nM labeled His-TREM2 was incubated with 100 µM compound in assay buffer (PBS, pH 7.4, 0.05% Tween-20, final DMSO concentration 2%) on Dianthus plates for 15 minutes. Additionally, a negative control (PBS, pH 7,4 0.05% Tween-20, 2% DMSO) was run in one column per plate as well as a positive control (100 µM, **T2337**).^21^ Before running the HTS setup, we determined a Z’ factor for the assay (**Table S1**). Additionally, we determined the Z’ value for each plate in the screening (**Figure S2**). We analyzed the normalized fluorescence (F_norm_) for all wells using GraphPad Prism 10. Compounds with an F_norm_ outside of a ten standard deviation range from the average control were considered as potential hits. Compounds that were flagged by the instrument’s software (DI.Control) for aggregation or scan anomalies were excluded from the analysis. A more detailed description of the hit selection can be found in the **Supporting Information**.

#### Control Experiments

We performed control experiments to exclude assay interfering compounds. Firstly, the hit compounds were incubated with assay buffer and the initial fluorescence and TRIC signal were assessed with Dianthus. For a second control, all non-interfering compounds from the first experiment were incubated with RED-tris-NTA 2^nd^ generation dye and their initial fluorescence and TRIC signal were measured using Dianthus. Compounds that did not show any significant difference from the average reference (PBS, pH 7.4, 0.05% Tween20, 2% DMSO) were considered for further experiments. All remaining hits were verified with a repetition of the single-dose screening at 100 µM in two additional independent experiments.

### Binding Affinity with Monolith X

All experiments were performed on a Monolith X (NanoTemper Technologies) using Monolith Premium Capillaries (NanoTemper Technologies, #MO-K025). Recombinant human His-tagged TREM2 (SinoBiological, #11084-H08H) was labeled with RED-tris-NTA 2^nd^ generation dye (NanoTemper Technologies, #MO-L018) according to the manufacturer’s protocol and the protein concentration was adjusted to 40 nM. The ligand solutions were prepared as 2-fold stocks at 1 mM or 800 µM in 2-fold assay buffer (PBS, pH 7.4, 0.05% Tween-20, 10% DMSO) and subsequently diluted 1:1 with assay buffer to get a 16-point dilution series with 10 µL for each sample. Then, all samples were incubated with 10 µL of a 2-fold protein stock (40 nM) in PCR tubes for 15 min at room temperature (final concentrations: 500–0.02 or 400–0.01 µM ligand, 20 nM His-TREM2, 5% DMSO). Capillaries were dipped into each tube, loaded onto the chip, and the measurement was started at 100% excitation with medium MST power. Thermophoresis was measured for 10 seconds with an initial five second delay. All samples were run in three independent experiments. The raw TRIC data were analyzed using the instrument’s software (MO.Control 2) to create merge sets of all replicates per compound (see **Supporting Information, Figures S7–9**). Capillaries that were flagged by the software as initial fluorescence outliers or for aggregation were excluded from the analysis. For better comparability between different compounds, the data was normalized to fraction bound.

### Selectivity Studies

#### LAG-3

All experiments were performed as previously published with a few adjustments.^22^ The samples were run on a Monolith X (NanoTemper Technologies) instrument. The screening was conducted at 300 µM ligand concentration, 2% DMSO. Thermophoresis was measured for 15 seconds with an initial five second delay. Data sets were analyzed using the instrument’s software (MO.Control 2). Single-dose screening results (F_norm_) were plotted and compared to a DMSO control (three standard deviation range from average).

#### TREM1

Binding affinity measurements were performed using microscale thermophoresis (MST) on a Monolith NT.115 system (NanoTemper Technologies). Recombinant human TREM1 protein (BioTechne, Minneapolis, MN, USA, #10337-TR) was labeled using the RED-tris-NTA 2^nd^ generation His-tag labeling kit (NanoTemper Technologies, #MO-L018) according to the manufacturer’s instructions. MST measurements were conducted using PBS buffer (pH 7.4) with 0.05% Tween-20. Labeled proteins were used at a final concentration of 40 nM and incubated with serially diluted test compounds for 10 minutes at room temperature (22–25°C) prior to measurement. MST experiments were performed using standard capillaries with the following instrument parameters: red filter set, 100% LED power, and medium MST power. Thermophoresis was monitored for 20 seconds with an additional 5-second delay. Data analysis was conducted using MO.Affinity Analysis software (NanoTemper Technologies).

### Thermal Shift Assay (TSA)

The experiments were run on a Prometheus Panta instrument (NanoTemper Technologies) using Prometheus High Sensitivity Capillaries (NanoTemper, #PR-C006). All buffers were filtered with 0.2 µm syringe filters before use. His-tagged TREM2 (SinoBiological, #11084-H08H) was reconstituted, and the concentration was adjusted to 20 µM (0.38 mg/mL) followed by a buffer exchange to PBS, pH 7.4, 0.05% Tween20 using Zeba™ Spin Desalting Columns for microcentrifuges (ThermoFisher Scientific, #89882). The protein was centrifuged for 10 min at 15,000 g, 4 °C before use. For the screening, ligand solutions were prepared as 2-fold stocks at 200 µM in assay buffer (PBS, pH 7.4, 0.05% Tween-20, 4% DMSO). Then, 12.5 µL of the ligand solutions were incubated with 12.5 µL of the protein stock in PCR tubes, resulting in 100 µM samples in 2% DMSO with 10 µM His-TREM2 (0.19 mg/mL). The samples were incubated for 10 min at room temperature before dipping capillaries into the tubes and starting the measurement.

The temperature gradient followed a 2 °C/min ramp from 30–85 °C and each sample was run in duplicates. Compound **T2337** was used as a positive control under the same conditions.^21^ The raw data were analyzed using the Panta analysis software (PR.Panta Analysis) followed by plotting the data in GraphPad Prism 10. For dose-dependent experiments, ligand solutions were prepared as 2-fold stocks at 400 µM in assay buffer (PBS, pH 7.4, 0.05% Tween-20, 4% DMSO) followed by 1:1 dilutions with assay buffer to get a 5-point dilution series. Then, all samples were incubated with the protein stock as described above and incubated at room temperature for 10 min. The final ligand concentrations were 200, 100, 50, 25 and 12.5 µM in PBST with 2% DMSO. Compound **T2337** was used as a positive control under the same conditions (**Figure S11**).^21^ The measurement and data analysis were conducted as described above with three independent experiments per compound. Data were plotted in GraphPad Prism 10 to determine EC_50_ values.

### AlphaLISA Assay for SYK and DAP12 Phosphorylation

Phospho-AlphaLISA assays were used to quantify the phosphorylation of SYK and DAP12 at specific tyrosine residues (Tyr525/526 for SYK and Tyr91 for DAP12). This assay employs two antibodies: one recognizing the phosphorylated epitope and another targeting a distinct epitope on the same protein. In the presence of the phosphorylated target, the proximity of donor and acceptor beads generates a luminescent Alpha signal, the intensity of which is directly proportional to the amount of phosphorylated protein in the sample (according to the manufacturer’s description, Revvity). Total SYK and DAP12 protein levels were measured using corresponding AlphaLISA Total assays for data normalization.

HEK-hTREM2/DAP12 and wild-type HEK (lacking TREM2 and DAP12 expression) cells were seeded at a density of 50,000 cells per well in 96-well plates, in a final volume of 100 μL of DMEM (Gibco) supplemented with 10% fetal bovine serum (FBS; Gibco). Cells were incubated for 24 h at 37 °C in a humidified 5% CO₂ atmosphere. Test compounds **VG-3927** and **6B10-9** were prepared in DMEM at a final concentration of 25 μM; DMSO was used as a vehicle control at the corresponding final concentration. Following serum starvation, the culture medium was removed and replaced with compound-containing medium. After 1 h of incubation, the medium was aspirated, cells were lysed, and the AlphaLISA assays were performed according to the manufacturer’s instructions (Revvity, USA).

### Phagocytosis Assay

Phagocytosis assay was performed using HMC3 human microglial cells overexpressing the TREM2 receptor. Fluorescent bioparticles conjugated with a pH-sensitive dye that emits fluorescence upon internalization were used at a final concentration of 1 mg/mL. HMC3 cells were seeded at a density of 1 × 10⁵ cells per well in 96-well plates in 100 µL of growth medium (EMEM, Gibco) supplemented with 10% FBS (Gibco) and incubated for 24 h at 37 °C in a humidified atmosphere with 5% CO₂. Following serum starvation, cells were treated with either compound **VG-3927**, **6B10-9** (25 µM for 30 min), or solubilization buffer (vehicle control) at the corresponding concentration. Conjugated bioparticles were then added to the medium and incubated for an additional 60 min. After incubation, cultures were washed three times with ice-cold PBS, and fluorescence intensity was measured using a Tecan Spark (Tecan Group, Austria) plate reader at excitation/emission wavelengths of 560/585 nm.

### In Silico Binding Analysis

The molecular docking employed in this study followed the same computational methodology as previously described in our recently published work.^23, 24^

## Results and Discussion

### High-Throughput Fragment Screening with Dianthus

Before screening, we determined a Z’ factor of 0.71 for the setup, which indicates a good suitability of our platform for HTS (see **Supporting Information**). The initial single-dose screening at 100 µM ligand concentration resulted in 50 potential hits (hit rate: 1.6%, **Figure 1A**). Here, we considered compounds that were outside of a ten standard deviation range from the average reference per plate (**Figure 1B**). By contrast, we did not consider compounds that were flagged by the Dianthus software for aggregation or scan anomalies (111 compounds, 3.5%). To exclude assay interfering compounds, we conducted control experiments without the protein (**Figure S3**), narrowing down the number of potential hits to 30 (hit rate: 0.94%, **Figure 1A**). Then, we tested the remaining compounds in two additional binding checks at 100 µM (**Figure S4**) to exclude compounds that gave a false positive signal in the first screening, resulting in a final number of 14 hits (hit rate: 0.44%, **Figure 1A**).

**Figure 1.**
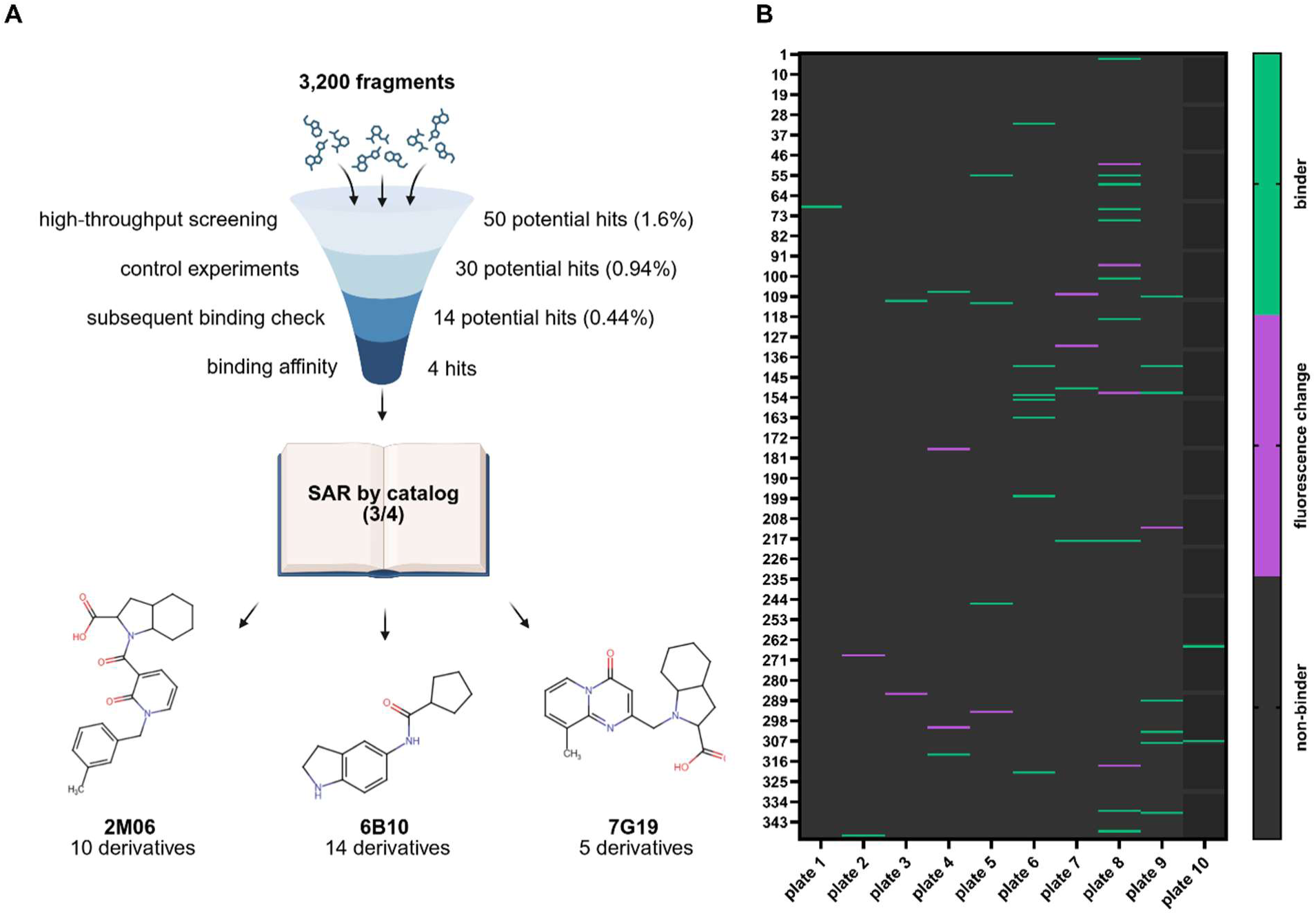
Overview of high-throughput fragment screening. (**A**) Workflow and yield for screening of PPI fragment library from initial HTS to hit derivatization for three out of four final compounds. (**B**) Screening results after single-dose HTS (100 µM ligand, 2% DMSO) for PPI fragment library. Green cells display binders within the same initial fluorescence range (±20%) as the reference (38 compounds), light purple cells represent potential binders that induced a change of the initial fluorescence (12 compounds). Plates 1–9: 352 compounds per plate (columns 1–22), plate 10: 32 compounds (columns 1+2). Hit map created with GraphPad Prism 10. Figure composed with BioRender.

### Hit Validation with Monolith

We purchased the remaining hit compounds for dose-dependent studies with Monolith X. Out of 14 compounds, three (**2M06**, **6B10**, **7G19**) could be validated as TREM2 binders with *K*_D_ values in the low to medium micromolar range (**Figure 2**, **2M06**: 26.1 µM, **6B10**: 37.6 µM, **7G19**: 3.83 µM). A fourth compound, **3N02**, also displayed dose-dependent binding in the higher micromolar range (*K*_D_ = 148 µM), but was not selected for follow up studies due to its lower potency compared to the top three. Overall, we were able to validate four out of 3,200 compounds (0.12%) as dose-dependent TREM2 binders.

**Figure 2.**
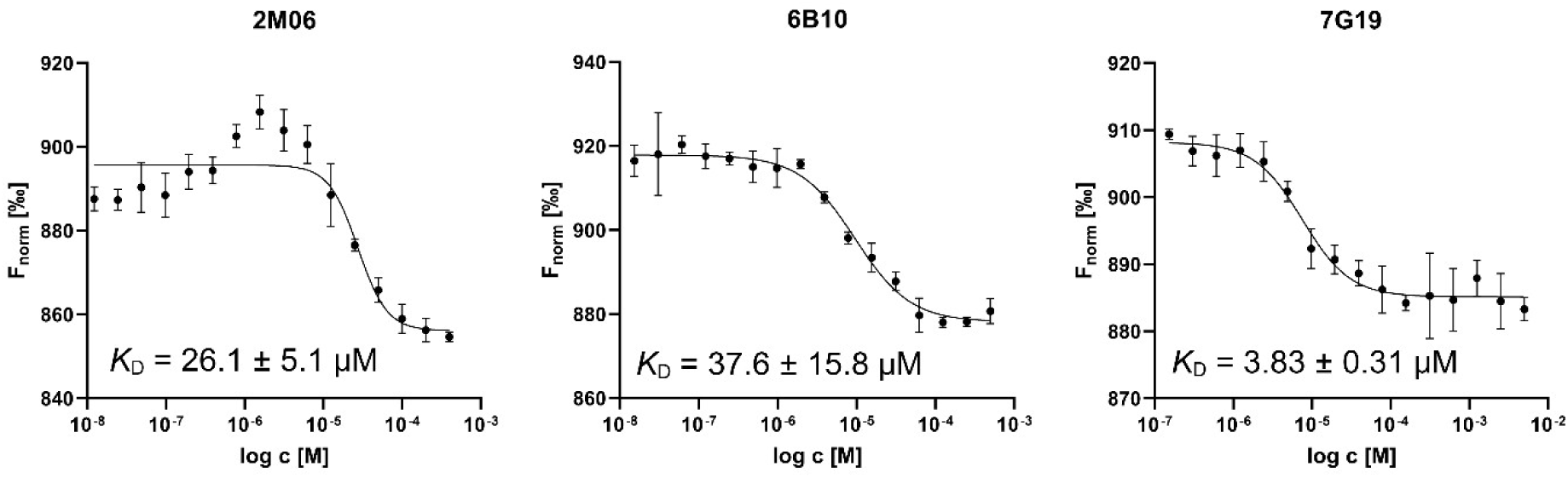
Hit validation using MST. Three compounds (**2M06**, **6B10**, **7G19**) emerged as dose-dependent binders of His-TREM2 with binding affinities in the low to medium micromolar range. Data are represented as means with standard deviations from three independent experiments. All graphs were created with GraphPad Prism 10.

### Hit Derivatization

#### SAR by catalog to find derivatives

After confirming the three top hits, we purchased several derivatives for each compound from Enamine to conduct a structure-activity relationship (SAR) study. We selected the derivatives based on modifications to different parts of the parent molecules. Compound **2M06** was divided into three major structural parts (**Figure 3**, top left corner), **6B10** and **7G19** into two parts (**Figure 3**, bottom and top right corner). In consideration of their individual availability and our expectations for the potency of the compounds, we purchased 10 derivatives for **2M06**, 14 for **6B10**, and 5 for **7G19**.

**Figure 3.**
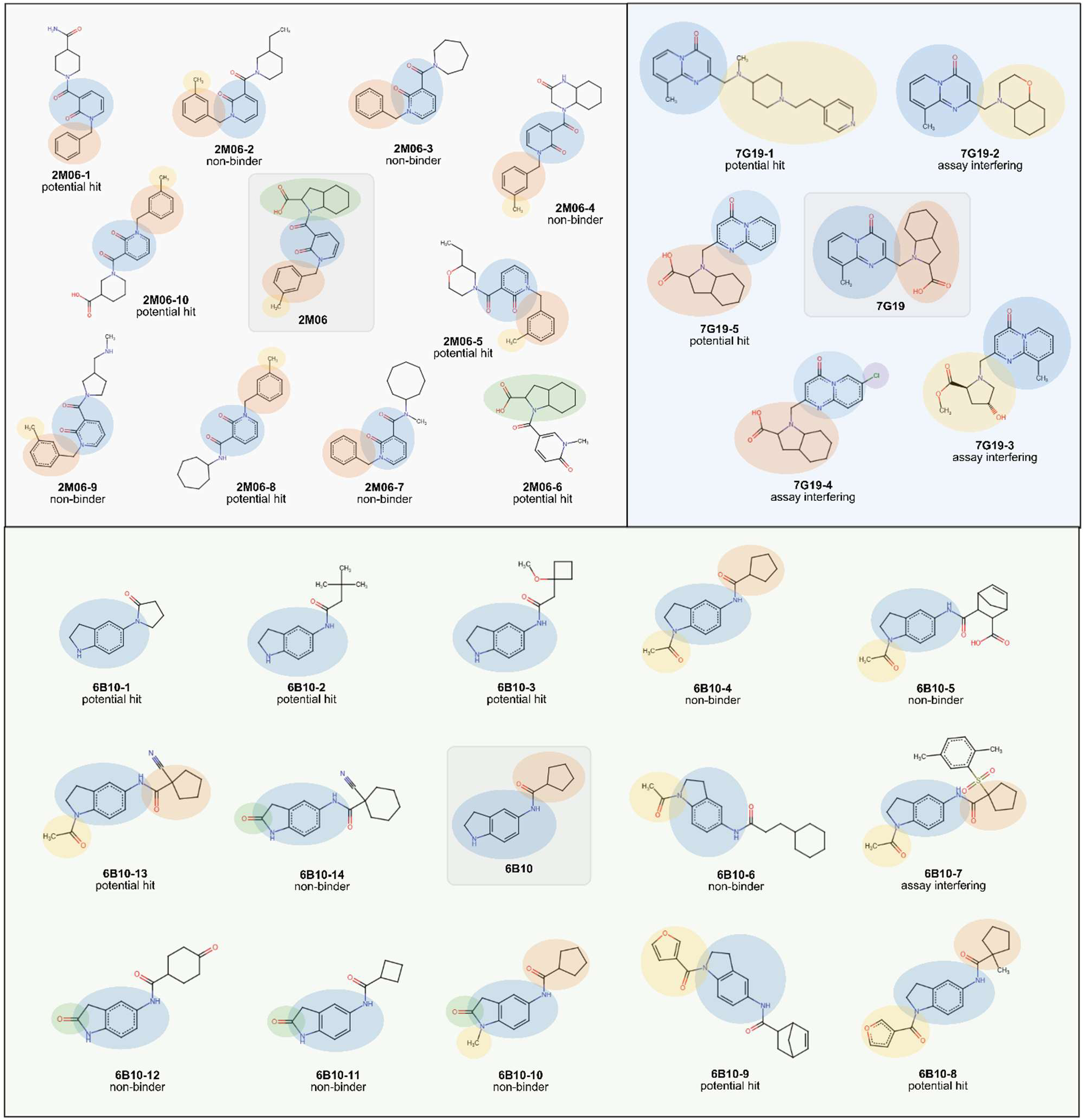
Results for hit derivatives after single-dose screening and control experiments. Top left corner: structures of **2M06** derivatives with assay results. Top right corner: structures of **7G19** derivatives including assay results. Bottom: structures of **6B10** derivatives with screening results. Figure composed with BioRender.

#### Initial Single-Dose Screening

Initially, we screened all 29 derivatives at a single dose of 100 µM with Dianthus as described above for the fragment library, followed by control experiments (**Figures S5** and **S6**). The experiments were conducted in three independent replicates and hits were considered based on the same standards as in the initial HTS. The screening identified five **2M06** derivatives (hit rate: 50%), six **6B10** derivatives (hit rate: 43%), and two **7G19** derivatives (hit rate: 40%) as potential hits (overall hit rate: 45%, see **Figure 3**). In the **6B10** subset, one compound (**6B10-7**, 7%) was interfering with the assay, whereas three compounds in the **7G19** subset (**7G19-2**, **7G19-3**, **7G19-4**, 60%) showed interference in the control experiments (**Figure 3**).

#### Hit Validation

We proceeded to evaluate the remaining 13 potential hit derivatives (**Figure 3**) in dose-dependent studies using Monolith X. Here, 11 out of the 13 showed dose-dependent binding of TREM2. However, five compounds (**2M06-6**, **2M06-8**, **6B10-2**, **6B10-3** and **6B10-13**) did not result in measurable *K*_D_ values since they did not reach plateaus at the highest ligand concentrations (**Figure 4A–C**). In the **2M06** series (**Figure 4A**), only derivative **2M06-1** resulted in a measurable *K*_D_ of 124 µM which does not outcompete its parent compound (*K*_D_ = 26.1 µM). For the **6B10** derivatives (**Figure 4B**), **6B10-1** (*K*_D_ = 19.5 µM) emerged as the most potent compound in the same range as the parent compound (*K*_D_ = 37.6 µM). Another notable derivative is **6B10-9** with a binding affinity in the medium micromolar range (*K*_D_ = 68.3 µM). In the **7G19** series (**Figure 4C**), derivative **7G19-5** (*K*_D_ = 4.5 µM) had a binding affinity in the low micromolar range in the same range as its parent compound (*K*_D_ = 3.8 µM). The other derivative, **7G19-1**, was less effective with a *K*_D_ value in the medium micromolar range (*K*_D_ = 70.1 µM). Out of all hits, **7G19** and its derivative **7G19-5** seem to be the most potent in terms of TREM2 binding affinity. A summary for all derivatives including structures and binding affinities can be found in the **Supporting Information**.

**Figure 4.**
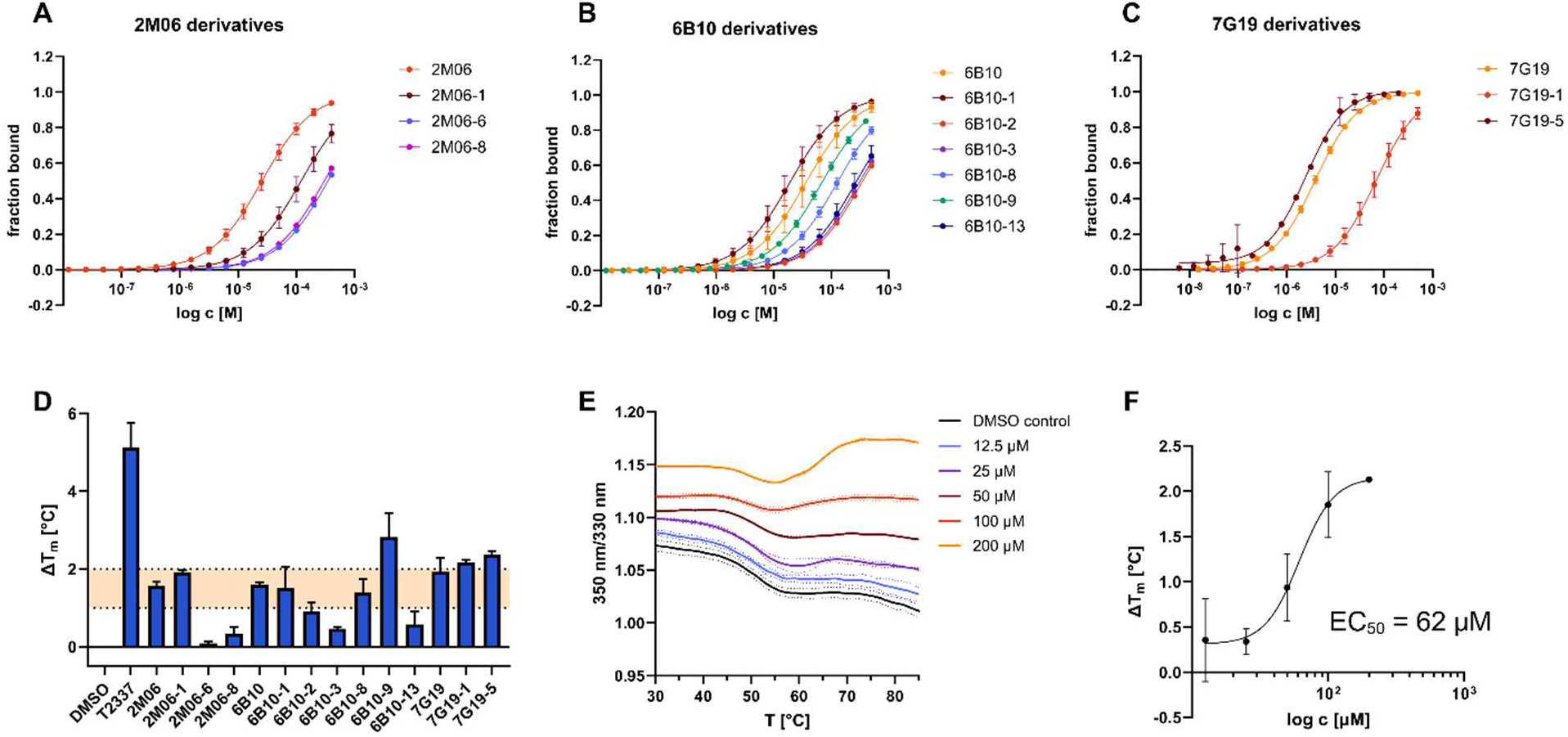
Overview of biophysical assays used to validate the derivatives. MST data represented as fraction bound graphs for better comparability, individual F_norm_ graphs in **Supporting Information** (**Figures S7–S9**). (**A**) **2M06** derivatives and their binding affinities with MST. *K*_D_ values in the low to medium micromolar range with the parent compound **2M06** (*K*_D_ = 26.1 µM) emerging as the set’s top compound. (**B**) **6B10** derivatives and their MST binding affinities in the medium micromolar range. Derivative **6B10-1** emerged as the most potent compound of the set (*K*_D_ = 19.5 µM). (**C**) **7G19** derivatives and their MST binding affinities. *K*_D_ values in the low to medium micromolar range with **7G19** (*K*_D_ = 3.8 µM) as the top hit of the set. (**D**) The average thermal shift in °C compared to the DMSO control (ΔT_m_) plotted as bar chart for all ligands. Data presented as means with standard deviations from three independent experiments. The ΔT_m_ range between 1 and 2 °C is highlighted in orange. **T2337** was used as a positive control. (**E**) NanoDSF-melting curves for His-TREM2 with different concentrations of ligand **6B10-9** including a DMSO control. Data presented as means with standard deviations from three independent experiments. (**F**) The average thermal shift in °C compared to the DMSO control (ΔT_m_) plotted against log c in µM for ligand **6B10-9**, resulting in a dose-response curve with an EC_50_ of 62 µM. All graphs were created with GraphPad Prism 10, figure composed with BioRender.

For additional validation of the original hits as well as their derivatives, we developed a thermal shift assay (TSA) for His-TREM2 based on nanoDSF. A detailed description and assay validation with a positive control can be found in the **Supporting Information** (**Figures S10** and **S11**). In contrast to differential scanning fluorimetry (DSF), nanoDSF is a label-free platform for melting curves that relies on the protein’s intrinsic tryptophan fluorescence.^25, 26^ The extracellular domain (ECD) of TREM2 (C-terminal His-tag, see materials and methods) includes four tryptophans, with two already being solvent-exposed, one buried, and one partially exposed.^5^ Upon melting, the protein unfolds which can be monitored in a change of fluorescence (ratio 350 nm/330 nm). After method optimization, we applied the technique to the initial validated hits and selected corresponding derivatives in a single-dose screening at 100 µM (**Figure 4D**, melting curves and control experiments in **Figure S12**). We observed that His-TREM2 (ECD) undergoes two unfolding events with a more pronounced transition at approximately 50 °C, which is in accordance with literature findings.^5^ A second inflection point can be noted at approximately 64 °C (**Figure S10**, **Figure 4E**), indicating that there might be another structural change along the temperature gradient. Due to the higher consistency regarding the first inflection point among independent experiments with His-TREM2 in buffer only, we examined the thermal shift around this melting temperature (50 °C) for our study. Here, only compound **6B10-9** induced a thermal shift of >2 °C compared to a DMSO control and was thus considered for dose-dependent experiments (**Figure 4E**). These studies suggested that **6B10-9** induces a dose-dependent thermal shift in the range of 200–12.5 µM compared to a DMSO control with an EC_50_ of around 62 µM (**Figure 4F**). This value is also reflected in the *K*_D_ value that was obtained with MST, 68.3 µM.

### Selectivity Studies

We selected the closely related triggering receptor on myeloid cells 1 (TREM1) and the unrelated immune receptor lymphocyte-activation gene 3 (LAG-3) as off-targets for selectivity studies. TREM1 shares a high structural similarity with TREM2 while playing an opposing role in the immune system.^2^ Both proteins mediate their effects through the adaptor protein DAP12, so it is crucial to determine the selectivity of the compounds before promising hits are taken forward into cell-based assays.^2^ TREM2 and LAG-3 are not structurally similar and belong to different families of immune receptors (TREM2 = TREM family,^2^ LAG-3 = CD4 homolog^27^), but both are involved in neurodegeneration (e.g., Alzheimer’s Disease) and cancer immunology.^1^ LAG-3 selectivity is important to identify potential non-specific binding.

To determine the selectivity, we conducted single-dose screenings at 300 µM using TRIC, then followed up with dose-dependent assays for potential hits (F_norm_ outside of three standard deviation range compared to DMSO control). We also presented the data as selectivity indices (SI) to enable better comparability, with higher values indicating a higher selectivity for TREM2 (**Table 1**). Most of the compounds for which we were able to determine TREM2 binding affinities displayed moderate to high selectivity against off-targets, e.g. **7G19** and **7G19-5** with at least 10-fold selectivity towards TREM1 and LAG-3. The **6B10** series was less selective but still exhibited a considerable 2–8-fold selectivity for TREM2. A limiting factor for the correct determination of selectivity indices was the lack of activity of most compounds in the single-dose screening for TREM1 and LAG-3 at a concentration of 300 µM, after which we did not conduct any further investigations into binding affinity. However, this suggests that most derivatives do not bind to these off-targets even a high ligand concentration, indicating specific binding to TREM2.

**Table 1.** Selectivity panel for selected derivatives. K_D_ values are determined by Monolith X, compared for TREM2 and off-targets, and expressed as selectivity indices (SI).

| cpd | $K_D$ (TREM2)<br>[$\mu\text{M}$] | $K_D$ (TREM1)<br>[$\mu\text{M}$] | SI (TREM2/<br>TREM1) | $K_D$ (LAG-3)<br>[$\mu\text{M}$] | SI (TREM2/<br>LAG-3) |
| --- | --- | --- | --- | --- | --- |
| 2M06 | $26.1 \pm 5.1$ | >300 | >11 | >300 | >11 |
| 2M06-1 | $124 \pm 36$ | >300 | >2.4 | >300 | >2.4 |
| 2M06-6 | >300 | >300 | $\approx 1$ | >300 | $\approx 1$ |
| 2M06-8 | >300 | >300 | $\approx 1$ | >300 | $\approx 1$ |
| 6B10 | $37.6 \pm 15.8$ | $66.9 \pm 28.6$ | 1.8 | >300 | >8.0 |
| 6B10-1 | $19.5 \pm 6.8$ | $121 \pm 3.4$ | 6.2 | $59.4 \pm 14.1$ | 3.0 |
| 6B10-2 | >300 | >300 | $\approx 1$ | >300 | $\approx 1$ |
| 6B10-3 | >300 | >300 | $\approx 1$ | >300 | $\approx 1$ |
| 6B10-8 | $127 \pm 16$ | >300 | >2.4 | >300 | >2.4 |
| 6B10-9 | $68.3 \pm 2.8$ | >300 | >4.4 | >300 | >4.4 |
| 6B10-13 | >300 | >300 | $\approx 1$ | >300 | $\approx 1$ |
| 7G19 | $3.8 \pm 0.3$ | $46.5 \pm 38.1$ | 12 | $61.9 \pm 23.2$ | 16 |
| 7G19-1 | $70.1 \pm 20.0$ | >300 | >4.3 | >300 | >4.3 |
| 7G19-5 | $4.5 \pm 0.9$ | $56.6 \pm 8.6$ | 13 | >300 | >67 |

### In Vitro Validation of 6B10-9-Mediated TREM2 Activation

Based on the previous results from MST and nanoDSF, compound **6B10-9** was further evaluated in a cell-based assay to assess its potential functional activity. To evaluate the agonistic effect of **6B10-9** on TREM2 signaling, we employed HEK293 cells that stably express human TREM2 and its signaling adaptor DAP12 (HEK293-hTREM2/DAP12). Phosphorylation of SYK and DAP12 was used as a proximal readout of TREM2 activation. To confirm the specificity of **6B10-9** activity, the compound was also tested in HEK293 cells lacking TREM2 and DAP12 expression. Treatment with **6B10-9** (25 µM, 1 h) induced robust phosphorylation of SYK and DAP12 in HEK293-hTREM2/DAP12 cells (**Figure 5B and 5C**). In contrast, no SYK phosphorylation was detected in cells lacking TREM2/DAP12, indicating that **6B10-9** specifically activates TREM2-dependent signaling (**Figure 5A**). We also tested more selected derivatives (see **Supporting Information**, **Figure S16**) but **6B10-9** was the only compound to induce a significant TREM2 activation.

**Figure 5.**
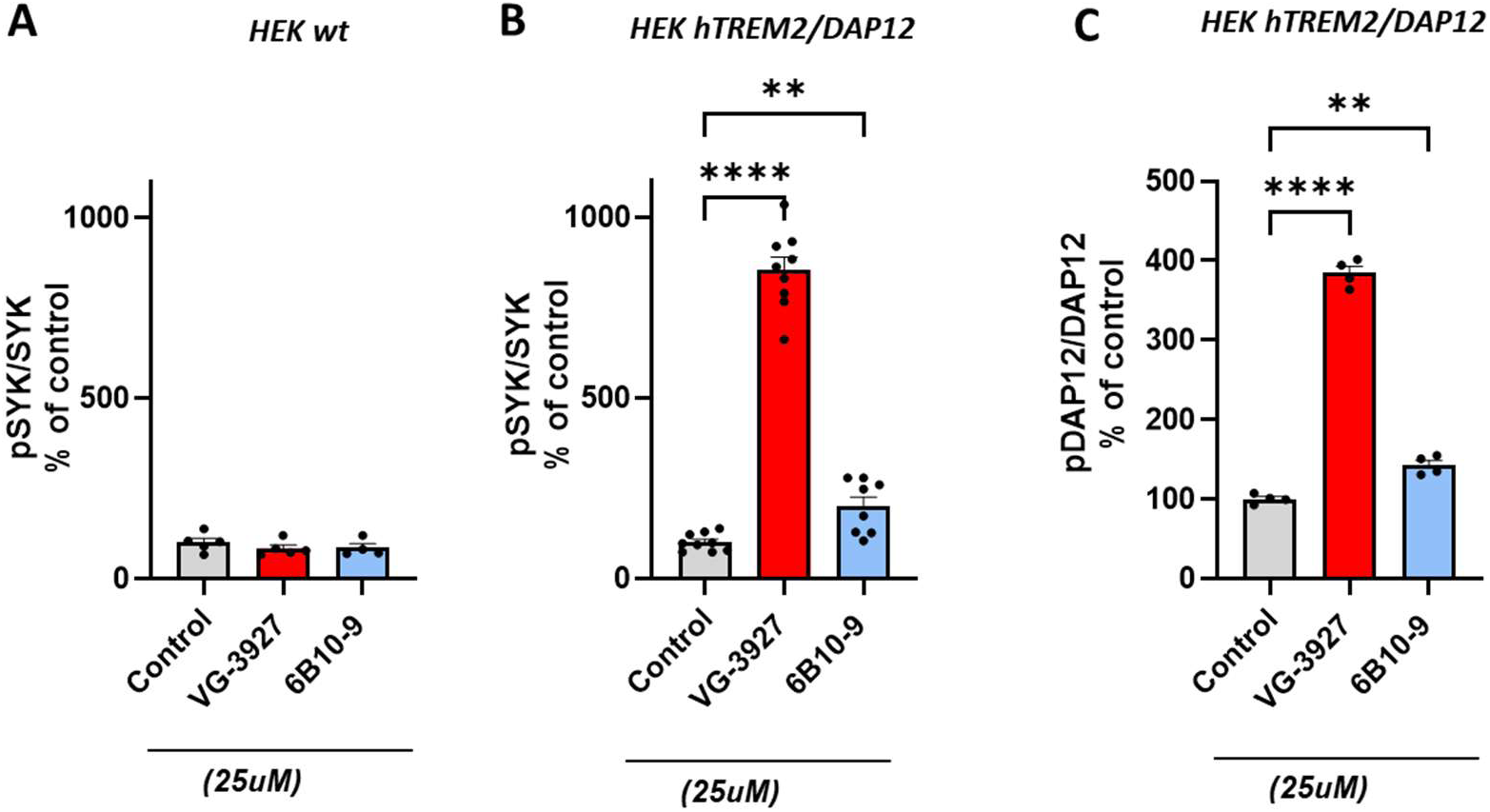
In vitro validation of the TREM2 binding activity. Histogram representing the quantification of phospho-SYK levels, measured using AlphaLisa technique, in untreated condition or after treatment with compound **VG-3927** (serving as reference) or **6B10-9** in non-expressing HEK cells **(A)** and in TREM2/DAP12 overexpressing HEK cells **(B)**. Cells were treated with 25 µM of compound for 1 h. Phospho-Syk changes in treated conditions are expressed as percentage of control **(B).** Histogram representing the quantification of phospho-DAP levels, measured using AlphaLisa technique, in untreated condition or after treatment with compound **VG-3927** or **6B10-9** in non-expressing HEK cells **(C)**. Data are presented as mean ± SEM from independent biological replicates (n = 4–8). Statistical analyses were conducted using Brown–Forsythe and Welch ANOVA tests, followed by Dunnett’s T3 multiple comparisons correction. Error bars represent means ± SEM. \*\**p* < 0.005, \*\*\*\**p* < 0.0001.

### 6B10-9 Enhances Microglial Phagocytosis

TREM2 activation is well known to promote microglial phagocytosis, a critical neuroprotective mechanism in neurodegenerative diseases.^28^ To determine whether **6B10-9** modulates this function, we examined its effect on phagocytic activity in human microglial HMC3 cells stably overexpressing human TREM2, while maintaining endogenous DAP12 expression. HMC3-hTREM2 cells were treated with either reference **VG-3927** or **6B10-9** (25 μM) for 30 minutes, followed by exposure to pHrodo™ bioparticles for 60 minutes. These bioparticles serve as phagocytic targets and emit fluorescence only upon internalization and acidification within phagolysosomes. The phagocytic activity was quantified by measuring the mean fluorescence intensity (MFI) of internalized pHrodo™ bioparticles. Data are presented as relative changes in MFI normalized to vehicle-treated controls. Treatment with **6B10-9** significantly increased pHrodo fluorescence intensity (70%) compared to vehicle-treated controls, indicating enhanced phagocytic uptake in TREM2-overexpressing human microglia cells (**Figure 6**). Interestingly, **7G19** and its derivatives also induced an increased phagocytic uptake (see **Supporting Information**, **Figure S15**) but had no significant effect in the phosphorylation assays.

**Figure 6.**
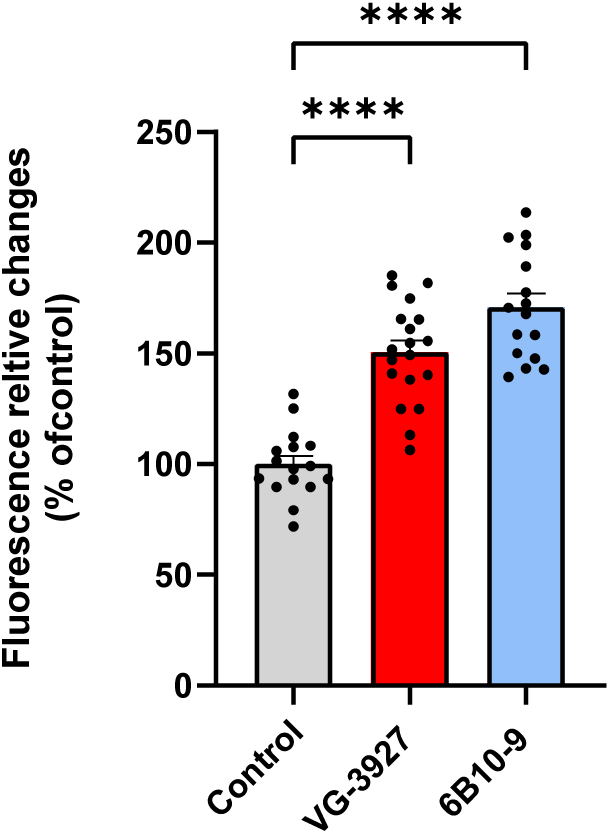
6B10-9 enhances phagocytic activity of human microglial cells. Phagocytic activity was quantified by measuring the mean fluorescence intensity of internalized pHrodo™ bioparticles using a plate reader. Mean fluorescence values for each condition were normalized to the control group and expressed as relative fluorescence change. HMC3-hTREM2 cells were treated with the indicated compounds (25 μM) for 30 min prior to exposure to pHrodo™ bioparticles for 60 min. **VG-3927** served as a reference, **6B10-9** increased phagocytic activity by approximately 70%. Data are presented as mean ± SEM from 15–19 biological replicates obtained across three independent experiments. Statistical analyses were performed using Brown–Forsythe and Welch ANOVA tests followed by Dunnett’s T3 multiple comparisons correction. ****p < 0.0001.

### Structure-Activity Relationship

In the **2M06** series, the parent compound emerged as the most potent (*K*_D_ = 26.1 µM). Additionally, one of the ten derivatives, **2M06-1**, proved to be a dose-dependent binder of TREM2 (*K*_D_ = 124 µM), but with more than 4-fold lower potency compared to **2M06**. In summary, only two out of eleven compounds in the series could be validated as dose-dependent TREM2 binders (19%), which does not provide reliable information on structure-activity relationships (SAR). However, the tested compounds displayed selectivity against off-targets like LAG-3 and the more closely related TREM1 in single-dose screenings. None of the hits from the series induced a significant effect in the thermal shift assay (ΔT_m_ <2 °C) or in cell-based assays.

By contrast, the **6B10** series provided more information on structure-activity relationships (**Figure 7**). Here, derivative **6B10-1** was most potent in terms of affinity (*K*_D_ = 19.5 µM), followed by parent compound **6B10** (*K*_D_ = 37.6 µM). In total, four out of fifteen (27%) showed dose-dependent TREM2 binding in MST with affinities in the medium micromolar range.

**Figure 7.**
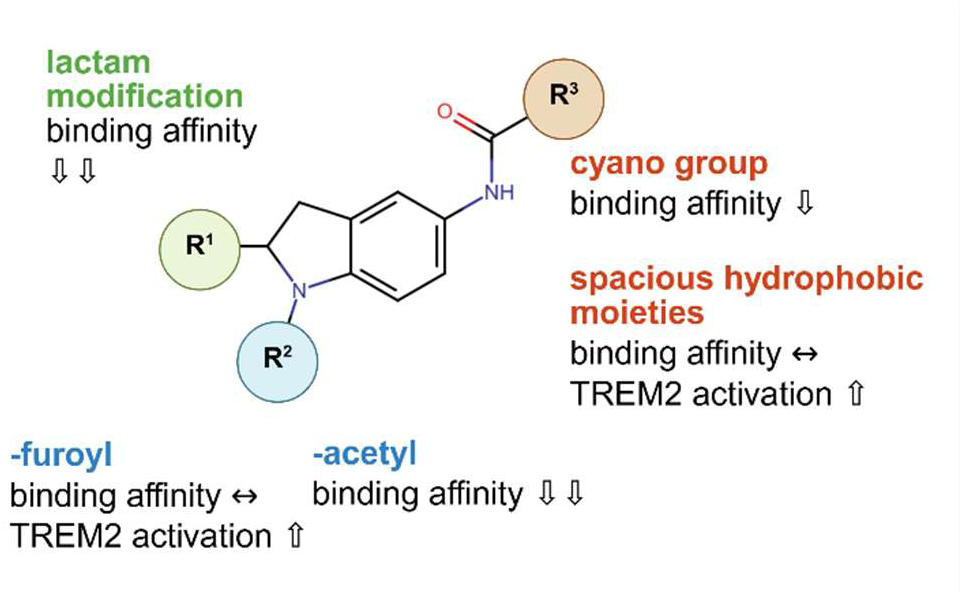
Structure-activity relationship for 6B10 derivatives. Results based on binding affinity and in vitro potency. Regarding binding affinity, *N*-acetylation of the indoline ring led to a loss of activity (**6B10-4** to **6B10-6**), whereas an addition of a furoyl group in the same position maintained TREM2 binding (**6B10-8** and **6B10-9**). Modifying the indoline ring with a lactam function also eliminated TREM2 binding (**6B10-10** to **6B10-12**). Additional modifications of the cycopentyl moiety only had a negligible effect on the potency compared to modifications at the indoline scaffold, e.g. the incorporated cyano group did not restore potency (**6B10-13** and **6B10-14**) after binding was abolished by alterations to the indoline moiety. In the thermal shift assay, all compounds that had a detectable *K*_D_ with TRIC induced a shift of 1–2 °C at 100 µM concentration, and compound **6B10-9** even resulted in a thermal shift of more than 2 °C, which we further validated to be dose-dependent (EC_50_ = 62 µM). In terms of selectivity, the **6B10** series was less selective than other derivatives. Compounds **6B10** and **6B10-1** share a high structural similarity, which is also reflected in their selectivity profiles. **6B10** was least selective in the set towards TREM1 (SI = 1.8) but more selective against LAG-3 (SI >8), whereas **6B10-1** was less selective towards LAG-3 (SI = 3). The structural modifications for **6B10-8** did not improve selectivity, which could partially be due to the higher *K*_D_ value for TREM2, since **6B10-9** with a binding affinity in a similar, but slightly higher, range as **6B10** and **6B10-1** displays a comparable selectivity profile. Therefore, the structural modifications did not lead to a significant improvement of selectivity. Nevertheless, the series shows a moderate selectivity towards off-targets which could be further improved by optimizing their binding affinity. In cell-based assays, **6B10-9** specifically activates TREM2-dependent signaling (DAP12 and SYK phosphorylation) and on top of that, significantly increases TREM2-mediated microglial phagocytosis. Parent compound **6B10** did not show any effects in these assays, indicating that the structural modifications of **6B10-9** (furanoyl group at indoline nitrogen and spacious hydrophobic moiety) could be essential in vitro. All in all, we successfully applied SAR by catalog to find more potent **6B10** derivatives in terms of biological activity which could serve as a good starting point for further optimizations towards a better binding affinity, and thus, higher selectivity.

The sample set of the **7G19** series was too small for reliable SAR conclusions. Here, three out of six compounds (**7G19-2** to **7G19-4**) interfered with the TRIC technology. However, the remaining three compounds all displayed dose-dependent TREM2 binding. The parent compound **7G19** as well as derivative **7G19-5** both have similar affinities in the low micromolar range (*K*_D_ = 3.8 and 4.5 µM, respectively). They share the same scaffold with only one minor difference in the methylation of the quinazolinone ring. These two compounds also exhibited activity against off-targets like TREM1 and LAG-3, with more than 10-fold selectivity for TREM2 in both cases. The other active derivative, **7G19-1**, has a more complex structure compared to **7G19** while maintaining the quinazolinone scaffold, which reduced the TREM2 binding potency (*K*_D_ = 70.1 µM) more than 18-fold, but did not display any activity against the off-targets in a single-dose screening. All three validated hits induced a thermal shift of approximately 2 °C at 100 µM in nanoDSF, but we did not perform dose-dependent studies to determine EC_50_ values. The compounds also enhanced phagocytosis in HMC3 cells but did not show a significant increase in TREM2-dependent phosphorylation of DAP12 and SYK in HEK cells. This indicates that further structural optimizations might be required to improve the performance in cell-based assays.

### In Silico Binding Study for 6B10-9

To investigate the binding mechanism of compound **6B10-9** with TREM2, molecular docking studies were performed using the crystal structure of TREM2 (PDB: 5ELI).^5^ The docking analysis revealed that **6B10-9** was predicted to occupy the same well-defined binding pocket within the TREM2 structure that we identified in our previous investigation (**Figure 8A**).^23^ The three-dimensional representation shows the protein adopting a characteristic immunoglobulin-like fold with multiple β-strands, creating a binding cavity suitable for ligand accommodation. The detailed interaction map (**Figure 8B**) suggested that the compound established several interactions with TREM2 binding site which consisted of a hydrogen bond with Asp86 as well as several π−π interactions to stabilize the predicted binding.

**Figure 8.**
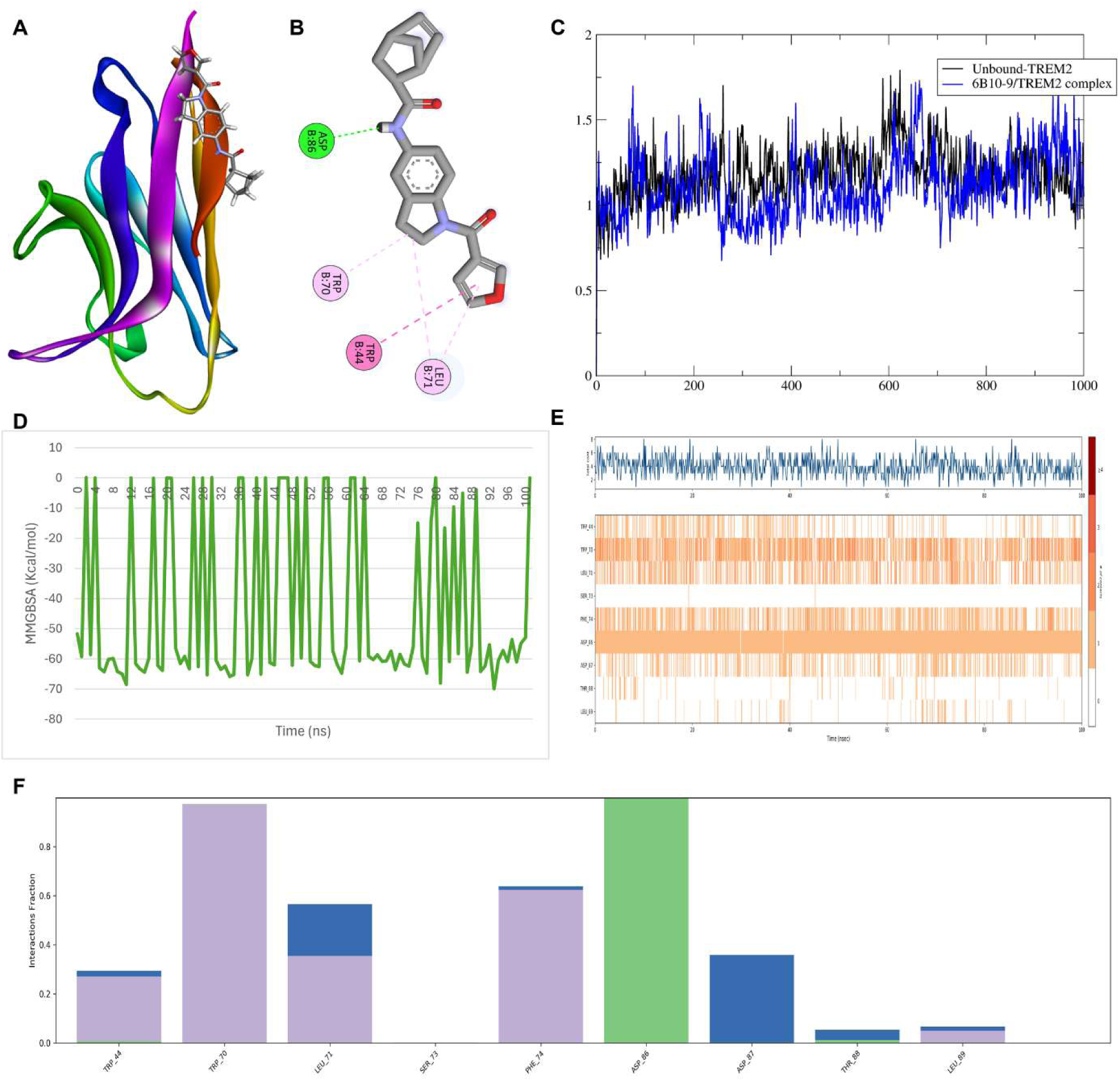
Molecular docking and dynamics simulation analysis of 6B10-9 binding to TREM2. **(A)** 3D structure of the TREM2 protein (PDB: 5ELI) showing the docked pose of compound **6B10-9**. (**B**) 2D interaction diagram depicting the binding mode of **6B10-9** with TREM2. (**C**) RMSD plot of Cα atoms as a function of simulation time (0-100 ns). (**D**) MM-GBSA analysis over 100 ns. (**E**) Time-dependent analysis of protein-ligand interactions over the simulation trajectory. Upper panel: Distance fluctuations between **6B10-9** and key TREM2 residues over time. Lower panel: Contact frequency heat map showing the probability of interactions between specific residue pairs throughout the simulation, with darker orange indicating higher contact occupancy. (**F**) Bond occupancy analysis showing the fraction of simulation time each bond was maintained. Green represents hydrogen bonds, purple represents π−π interaction while blue represents salt bridges.

To assess the stability and dynamic behavior of the **6B10-9**/TREM2 complex, molecular dynamics simulations were conducted over a 100 ns trajectory. The root mean square deviation (RMSD) analysis (**Figure 8C**) demonstrates the conformational stability of both the complex (blue line) and unbound TREM2 (black line) throughout the 100 ns of the MD simulation. Both systems exhibited RMSD values fluctuating between 0.5 and 1.8 Å, with the complex showing slightly higher flexibility in certain regions. The overall convergence of RMSD values indicates that the protein-ligand complex achieved equilibration and maintained structural integrity during the simulation, validating the stability of the predicted binding mode. The binding free energy landscape was further characterized through MM-GBSA calculations (**Figure 8D**), yielding an average binding energy of −43.60 kcal/mol. Structural dynamics were examined through contact analysis and secondary structure monitoring over the simulation trajectory (**Figure 8E**). The distance fluctuations between the ligand and key protein residues revealed persistent contacts maintained throughout the simulation. The contact analysis diagram indicated a stable and consistent contact across the entire 100 ns, where the hydrogen bond with Asp86 was maintained throughout the whole simulation. Similarly, the π−π interaction was maintained throughout the simulation which suggests its importance for stabilizing the binding of **6B10-9** with TREM2. This pattern of sustained interactions corroborates the RMSD findings and supports the formation of a thermodynamically stable complex. Finally, the hydrogen bond occupancy analysis (**Figure 8F**) predicted the relative contribution of specific interactions to complex stability. Multiple hydrogen bonds were identified, with varying occupancy fractions ranging from near-complete persistence (>0.8) to transient contacts (<0.3). The presence of high-occupancy hydrogen bonds, particularly involving residues such as Asp86 (green bar), indicates the formation of robust, persistent interactions that anchor the ligand within the binding pocket. Collectively, these computational analyses suggest that **6B10-9** forms a stable and energetically favorable complex with TREM2, characterized by persistent hydrogen bonding, favorable binding free energy, and maintained structural integrity throughout extended molecular dynamics simulations.

## Conclusions and Perspectives

In this study, we successfully applied our TRIC-based HTS platform to a fragment library of 3,200 compounds and identified new small molecule-based scaffolds for TREM2. We validated four fragment hits in dose-dependent studies with binding affinities in the micromolar range and then followed up with a “SAR by catalog” approach for three out of the four (**2M06**, **6B10**, **7G19**). Here, compound **6B10-9** emerged as the most promising candidate based on the combined results of microscale thermophoresis (*K*_D_ = 68.3 µM), a thermal shift assay (EC_50_ = 62 µM), selectivity profiles against TREM1 and LAG-3, and activity in cell-based assays (SYK and DAP12 phosphorylation and phagocytosis assay). Notably, several hits from the **7G19** series (**7G19-1**, **7G19-5**) also showed potential for follow up studies due to their binding potency in the low micromolar range, high selectivity, and an effect in microglial phagocytosis assays (see **Supporting Information**, **Figure S15**). Nevertheless, **6B10-9** demonstrated superiority in cell-based phosphorylation assays. A computer-aided binding analysis demonstrated that **6B10-9** forms a stable and energetically favorable complex with TREM2 through the formation of hydrogen bonds with residues such as Asp86. This complex remained stable throughout extended molecular dynamics simulations, indicating that the compound is robustly anchored in the binding site.

In summary, compound **6B10-9** can serve as a promising lead scaffold for TREM2-binding small molecules. Future optimizations could include a SAR expansion to improve the binding affinity from the mid-micromolar to a low- or sub-micromolar range. This may help to enhance the biological effect on SYK and DAP12 phosphorylation as well as microglial phagocytosis and could optimize the selectivity towards off-targets. Future directions could also include in vitro evaluation under disease-relevant conditions for neurodegeneration (e.g., models with amyloid plaques) and other preclinical studies like assessing pharmacokinetic properties (e.g., brain penetration and plasma stability).

## Acknowledgements

We acknowledge funding from the National Institutes on Aging (NIA) RF1AG084635 (PI: Gabr). We would also like to thank the Fisher Drug Discovery Resource Center of Rockefeller University (RRID:SCR_020985) for providing access to the NanoTemper Dianthus NT.23 Pico and NanoTemper Prometheus Panta.

## Abbreviations

DAP12: DNAX-activating protein of 12 kDa
DEL: DNA-encoded library
DSF: differential scanning fluorimetry
ECD: extracellular domain FBDD fragment-based drug discovery
HTS: high-throughput screening
LAG-3: lymphocyte-activation gene 3
MD: molecular dynamics
MST: microscale thermophoresis
RMSD: root mean square deviation
SAR: structure-activity relationship
SI: selectivity index
TAM: tumor-associated macrophage
TBDD: target-based drug discovery
TME: tumor microenvironment
TREM1: triggering receptor expressed on myeloid cells 1
TREM2: triggering receptor expressed on myeloid cells 2
TRIC: temperature-related intensity change
TSA: thermal shift assay
VS: virtual screening.

