## Supporting Information for "High-Throughput Fragment Screening Identifies a New Small Molecule Scaffold that Modulates TREM2 Signaling"

[**Figure S11.** Melting profile of His-tagged TREM2 in complex with positive control **T2337**. (**A**) Melting curve of His-TREM2 with **T2337** at different concentrations. Corresponding melting temperatures (T_m_) are indicated as dotted lines. (**B**) Melting temperature plotted against compound concentration to determine EC_50_ value. (**C**) For better comparability between replicates, the difference in T_m_ compared to the DMSO control is plotted against the compound concentration. All graphs were created from raw data with GraphPad Prism 10 and depict the average from three dependent experiments.	18](#_Toc214019502)

[**Figure S15.** Phagocytic activity in HMC3 cells, Histogram representing fluorescence intensity changes, reflecting internalization of pHrodo™ bioparticles. Mean fluorescence values for each condition were normalized to the control group and expressed as relative fluorescence change. HMC3-hTREM2 cells were treated with the indicated compounds (25 μM) for 30 min prior to exposure to pHrodo™ bioparticles for 60 min. **VG-3927** served as a reference, **7G19, 7G19-1** and **7G19-5** display increased phagocytic activity. Data are presented as mean ± SEM from 15 biological replicates. Statistical analyses were performed using Brown–Forsythe and Welch ANOVA tests followed by Dunnett’s T3 multiple comparisons correction. ****p < 0.0001.	21](#_Toc214019506)

[**Figure S16.** Histogram representing the quantification of phospho-DAP **(A)** and phospho-SYK **(B)** levels, measured using AlphaLisa technique, in untreated condition or after treatment with compound **VG-3927** (serving as reference) or **2M06, 7G19, 7G19-1** or **7G19-5** in TREM2/DAP12 overexpressing HEK cells. Cells were treated with 25 µM of compound for 1 h. Phospho-Syk and phospho-DAP changes in treated conditions are expressed as percentage of control. Data are presented as mean ± SEM from independent biological replicates (n = 4–8). Statistical analyses were conducted using Brown–Forsythe and Welch ANOVA tests, followed by Dunnett’s T3 multiple comparisons correction. Error bars represent means ± SEM. ***p* < 0.005, *****p* < 0.0001.	21](#_Toc214019507)

### High-throughput screening with Dianthus

#### Z’ factor determination

**Table S1.** Results for Z’ factor using **T2337** (100 µM, 2% DMSO) as a positive (purple) and 2% DMSO in PBST as a negative control (yellow).

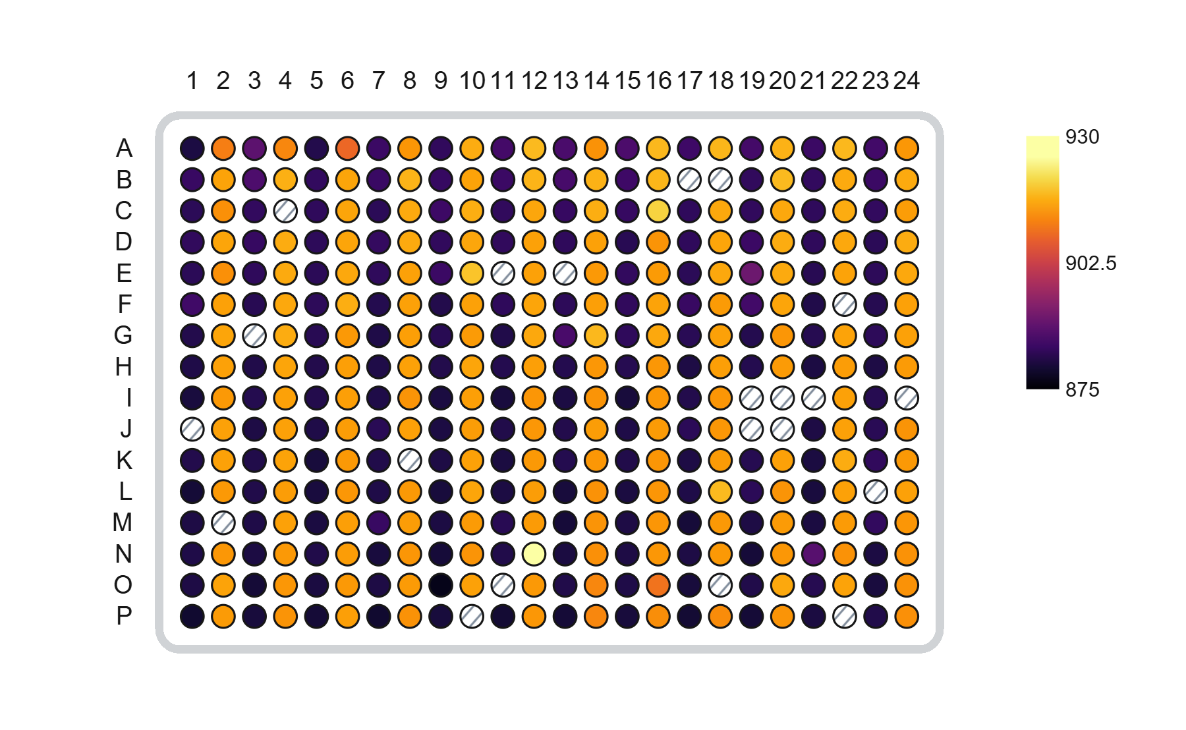

| avg. | 882.3 | 917.9 | 883.0 | 918.5 | 882.2 | 917.9 | 882.7 | 918.3 | 882.2 | 918.8 | 882.7 | 919.3 | 883.0 | 917.8 | 882.9 | 918.1 | 882.5 | 918.5 | 883.5 | 918.7 | 882.9 | 918.9 | 883.2 | 918.0 |
| --- | --- | --- | --- | --- | --- | --- | --- | --- | --- | --- | --- | --- | --- | --- | --- | --- | --- | --- | --- | --- | --- | --- | --- | --- |
| stdev | 1.65 | 1.26 | 2.53 | 1.27 | 1.20 | 1.88 | 1.57 | 1.05 | 2.08 | 1.24 | 1.74 | 3.08 | 2.35 | 1.64 | 1.89 | 2.44 | 1.60 | 1.40 | 3.00 | 1.32 | 2.09 | 1.04 | 1.42 | 1.09 |
| stdev [%] | 0.19 | 0.14 | 0.29 | 0.14 | 0.14 | 0.21 | 0.18 | 0.11 | 0.24 | 0.14 | 0.20 | 0.34 | 0.27 | 0.18 | 0.21 | 0.27 | 0.18 | 0.15 | 0.34 | 0.14 | 0.24 | 0.11 | 0.16 | 0.12 |
| Z' (per column pair) | 0.7544 | | 0.6789 | | 0.7412 | | 0.7794 | | 0.7280 | | 0.6041 | | 0.6550 | | 0.6316 | | 0.7500 | | 0.6309 | | 0.7390 | | 0.7830 | |
| **Av. Z’** | **0.7063** | | | | | | | | | | | | | | | | | | | | | | | |

To determine the Z’ factor, a known binder (**T2337**)^1^ was selected as a positive control (shown in yellow cells) and run on a 384-well Dianthus plate in alternating columns with the reference (PBST, 2% DMSO). The experiment was done under the same conditions as for the single-dose HTS screening. Blank cells indicate a scan anomaly or aggregation which occurred in less than 5% of all wells, potentially due to pipetting errors. Then, Z’ factors were calculated for each column pair, followed by the average Z’ out of all (**Table S1**). For this assay setup the overall Z’ is 0.71, confirming the suitability for HTS.

#### Plate layout

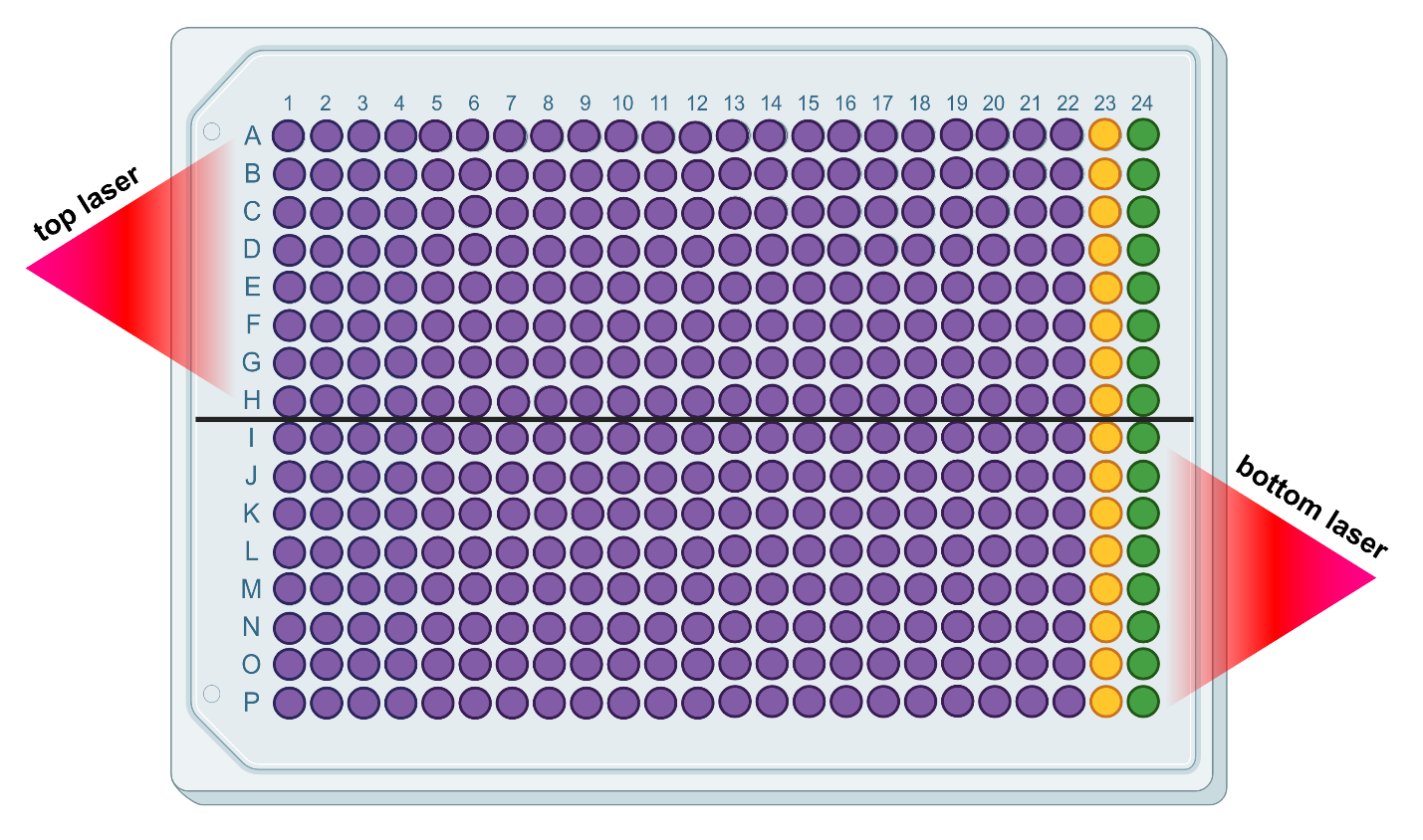

**Figure S1.** Plate layout including the location of the two lasers for plates 1–9. Plate 10: Compound library samples in columns 1–2, positive control in column 3, reference in column 4. Green wells represent the references (DMSO in PBST), yellow wells the positive control, and purple wells the samples. The top laser read wells A1–H24, the bottom laser wells I1–P24. Figure created with BioRender.

#### Hit selection

##### Initial single-dose screening

The results of the primary single-dose screening at 100 µM for each plate (3,200 compounds total) are shown in **Figure S2**. Compounds outside of a ten standard deviation range from the average reference on each plate were considered as potential hits. Compounds that were flagged by the Dianthus software as potential aggregators or that showed scan anomalies were excluded. The 50 initial hits were then tested in control experiments to exclude assay interfering false positives.

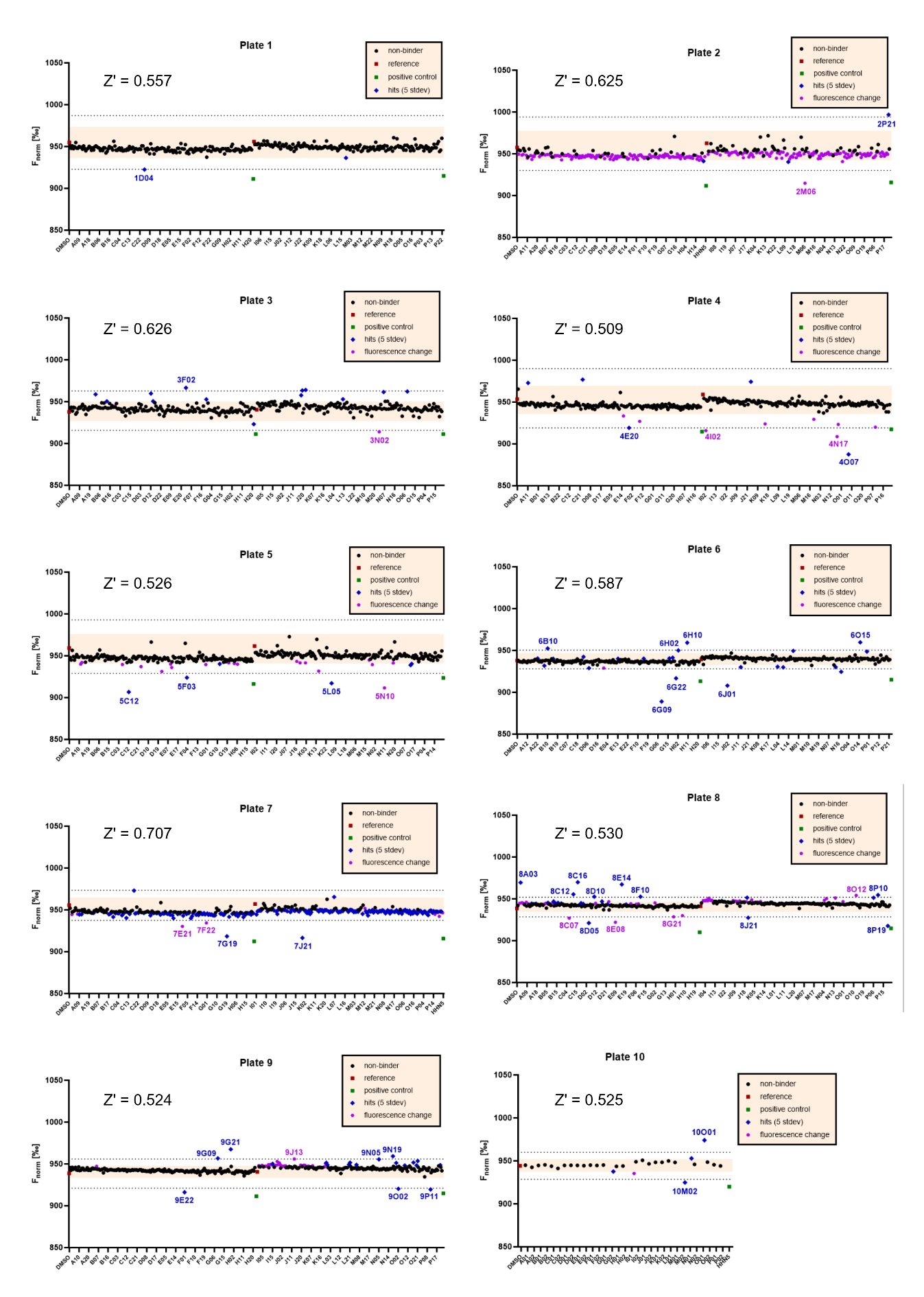

**Figure S2.** HTS results for plates 1–10. Compounds outside of a 10 standard deviation range from the average reference were considered as hits. Z’ values were calculated for each plate and added. Graphs created using GraphPad Prism 10, figure composed with BioRender.

##### Control experiments

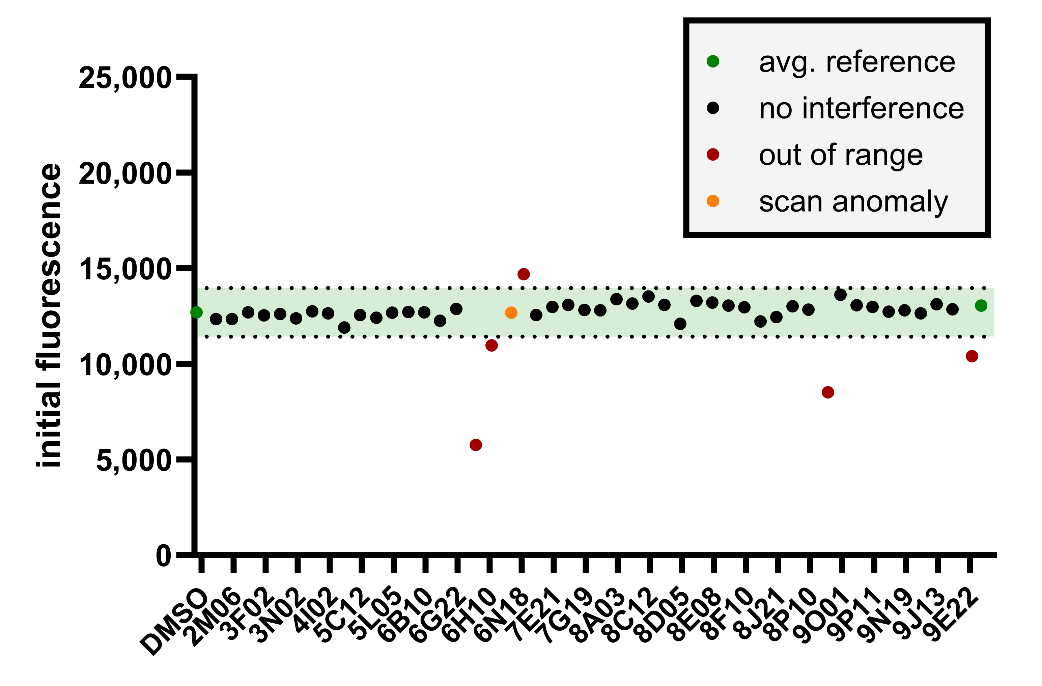

**Figure S3.** Results of control experiment. Average reference (n = 16) shown in green with a 20% range (light green) for initial fluorescence. Compounds within the range are depicted as black dots, out of range as red dots. Compounds within the range that showed scan anomalies are displayed as orange dots. Graph created with GraphPad Prism 10.

##### Additional binding check

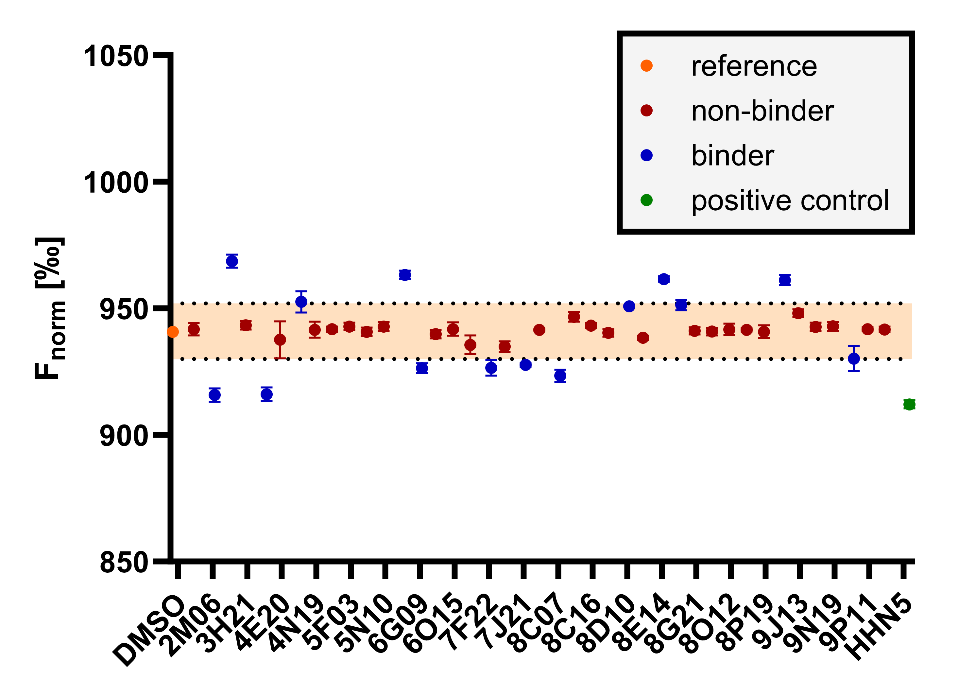

**Figure S4.** Results of two additional single-dose screenings at 100 µM, 2% DMSO. Average reference (DMSO control, n = 16) shown as orange dot with a ten standard deviation range (light red) for the F_norm_. Non-binders are depicted as red dots, binders outside of the range as blue dots. The positive control **T2337** (n = 16) is shown as a green dot for reference. Graph created with GraphPad Prism 10.

#### List of hit compounds

**Table S2.** List of hit compounds after control experiments and additional binding check at 100 µM.

| **well** | **cpd ID** | **MW [g/mol]** | **structure** | ***K*_D_ [µM]** |
| --- | --- | --- | --- | --- |
| 2M06 | Z1443635422 | 394.46 | 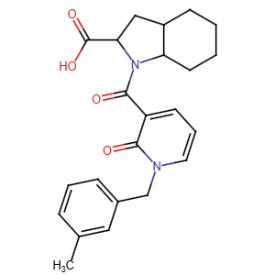 | 26.1 ± 5.1 |
| 3F02 | Z2418537157 | 255.74 | 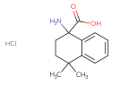 | n. d. |
| 3N02 | Z1222283881 | 279.21 | 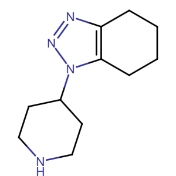 | 148 ± 21 |
| 4E20 | Z319921192 | 287.34 | 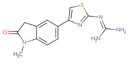 | n. d. |
| 4I02 | Z3303617233 | 300.80 | 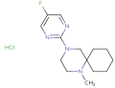 | n. d. |
| 6B10 | Z1456536982 | 266.77 | 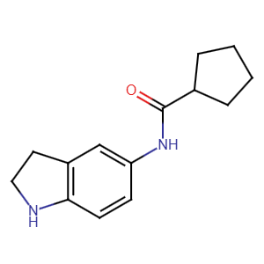 | 37.6 ± 15.8 |
| 6G09 | Z981544982 | 379.43 | 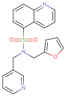 | n. d. |
| 7G19 | Z1276736199 | 341.40 | 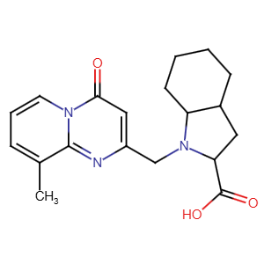 | 3.8 ± 0.3 |
| 7J21 | Z1348404668 | 287.36 | 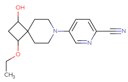 | n. d. |
| 8D10 | Z1724204436 | 316.44 | 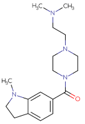 | n. d. |
| 8E14 | Z1457729446 | 348.82 | 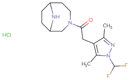 | n. d. |
| 8F10 | Z970557626 | 313.42 | 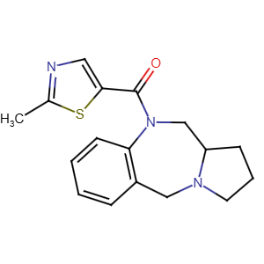 | n. d. |
| 9G21 | Z1155911897 | 335.45 | 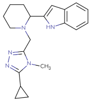 | n. d. |
| 9O02 | Z46159109 | 284.36 | 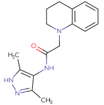 | n. d. |

n. d. not determined.

### Hit derivatization

#### Single-dose screening

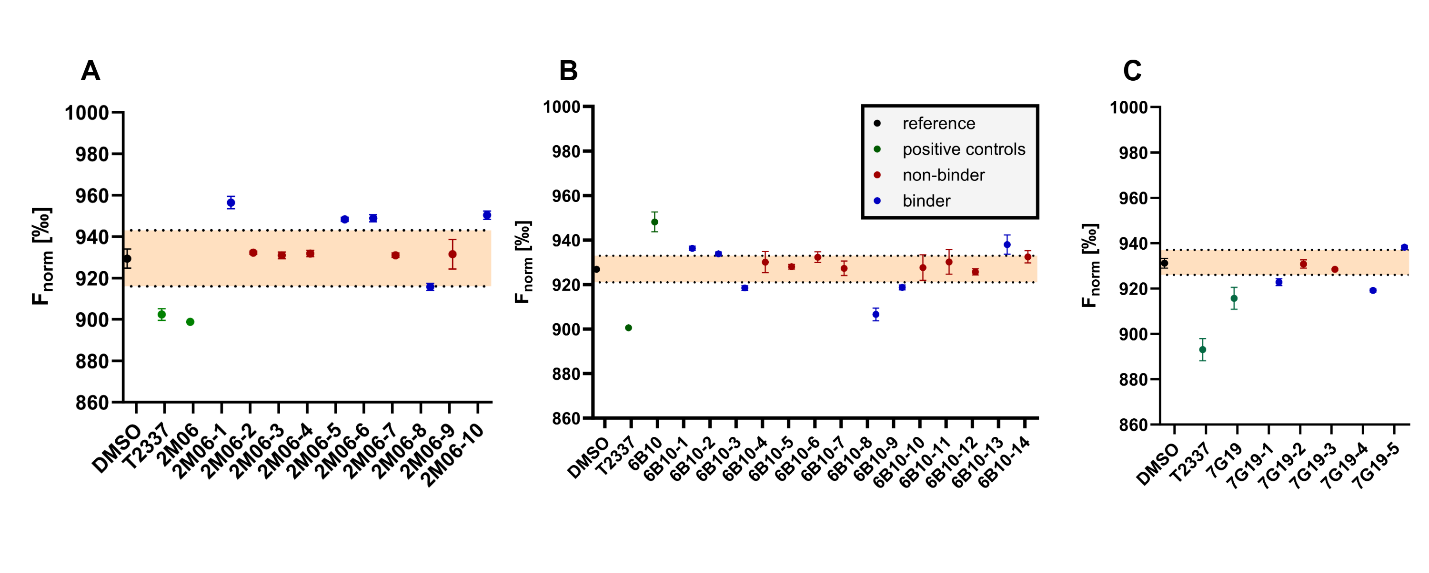

**Figure S5.** Results of single-dose screenings at 100 µM, 2% DMSO. Average reference (DMSO control, n = 6) shown as orange dot with a ten standard deviation range (light red) for the F_norm_. Non-binders are depicted as red dots, binders outside of the range as blue dots. The positive control **T2337** (n = 6) in assay buffer is shown as a green dot for reference. Graph created with GraphPad Prism 10.

#### Control experiments

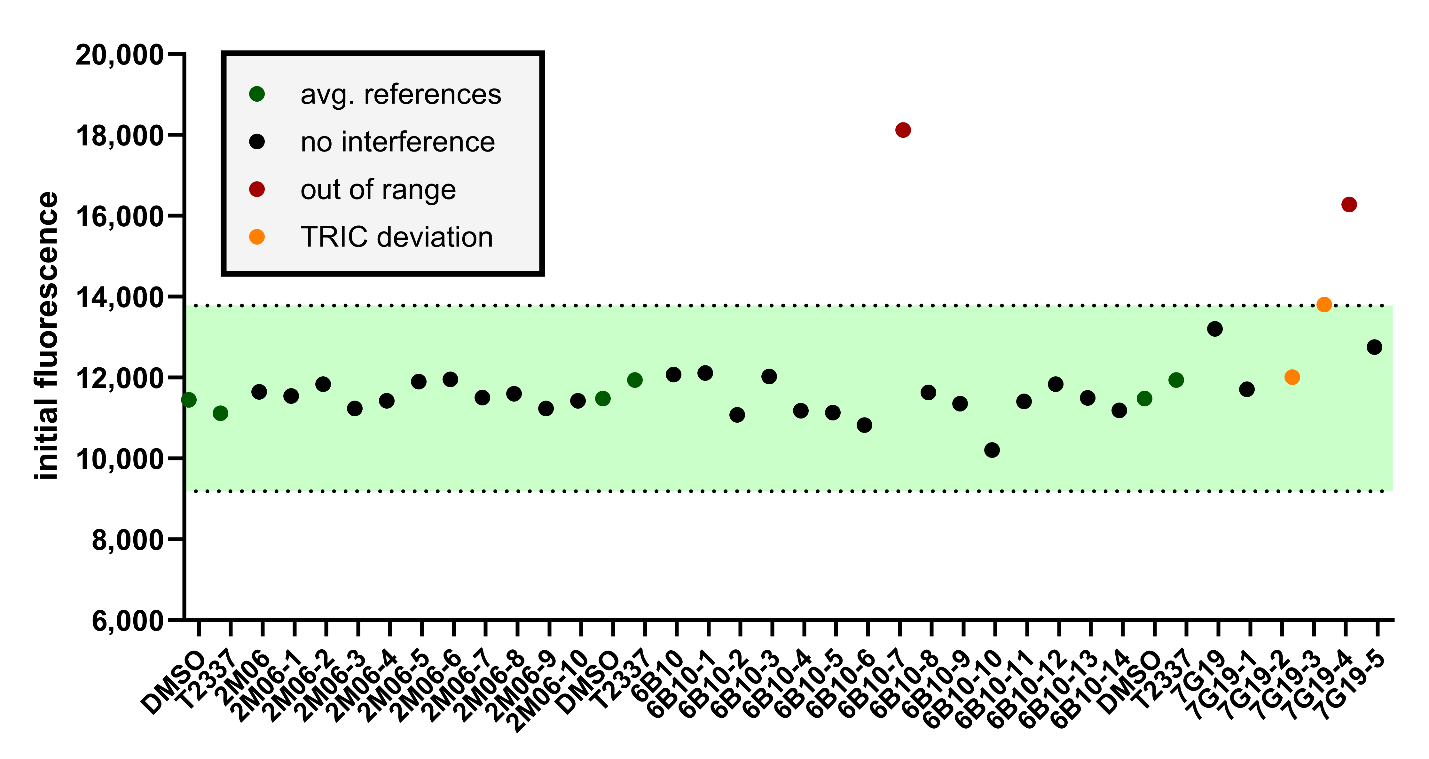

**Figure S6.** Average reference (n = 12) shown in green with a 20% range (light green) for initial fluorescence. Compounds within the range are depicted as black dots, out of range as red dots. Compounds within the range that showed TRIC trace deviations are displayed as orange dots. Graph created with GraphPad Prism 10.

#### Binding affinity with MST

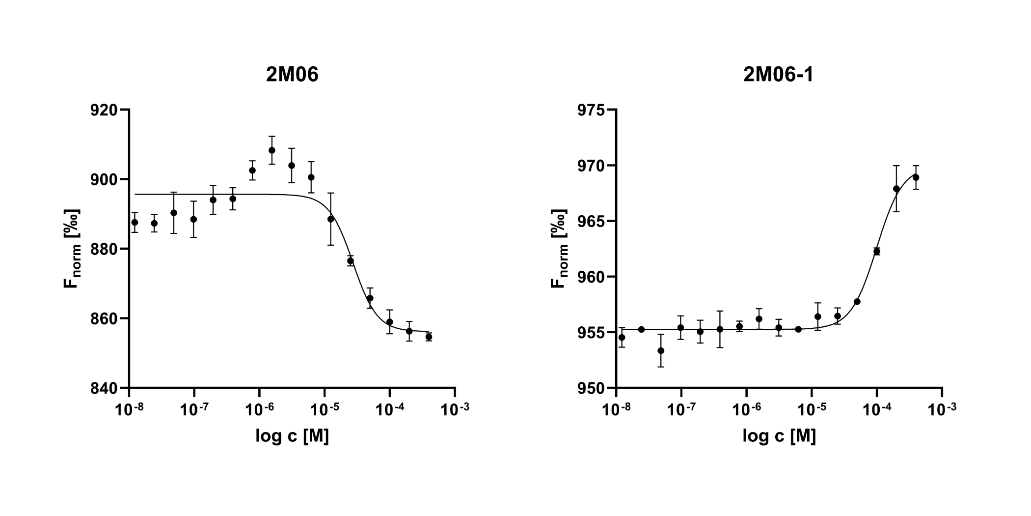

**Figure S7.** F_norm_ vs log c plots for **2M06** derivatives from three independent experiments per compound. Graphs created with GraphPad Prism 10.

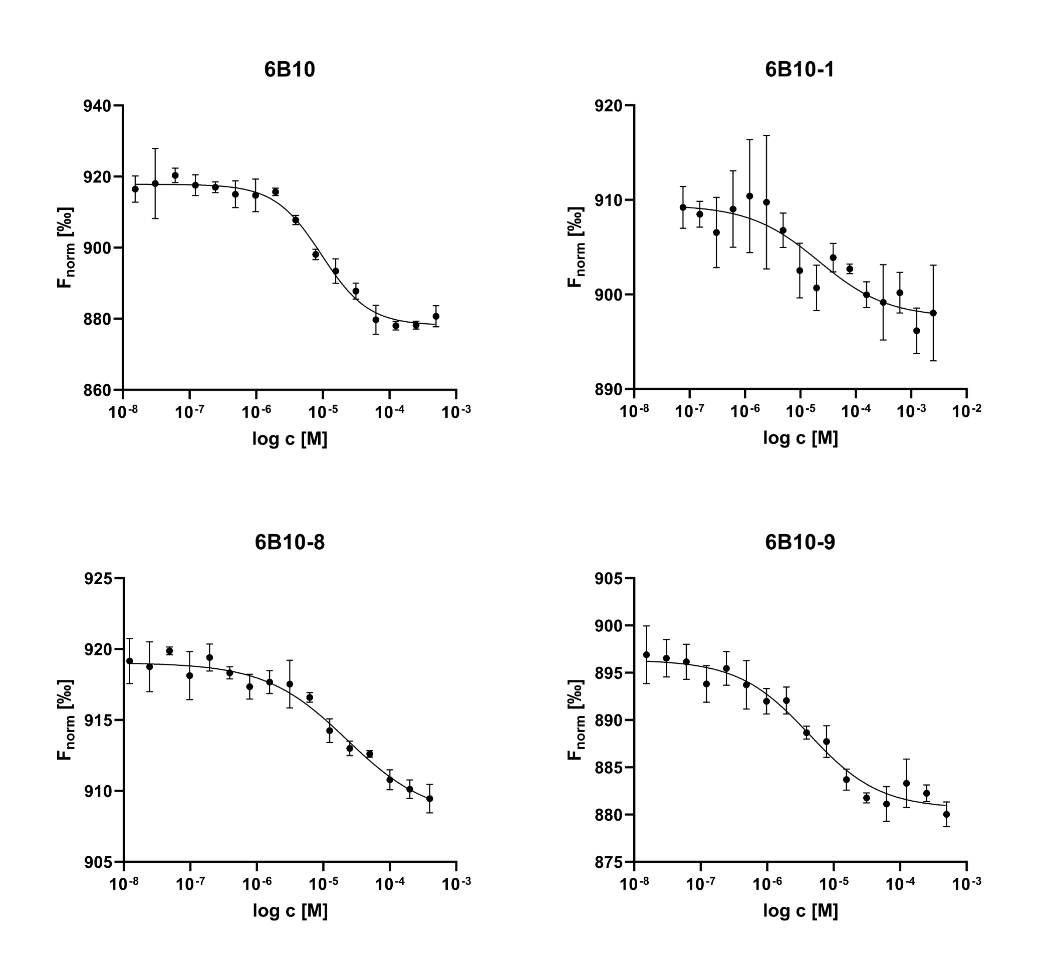

**Figure S8.** F_norm_ vs log c plots for **6B10** derivatives from three independent experiments per compound. Graphs created with GraphPad Prism 10.

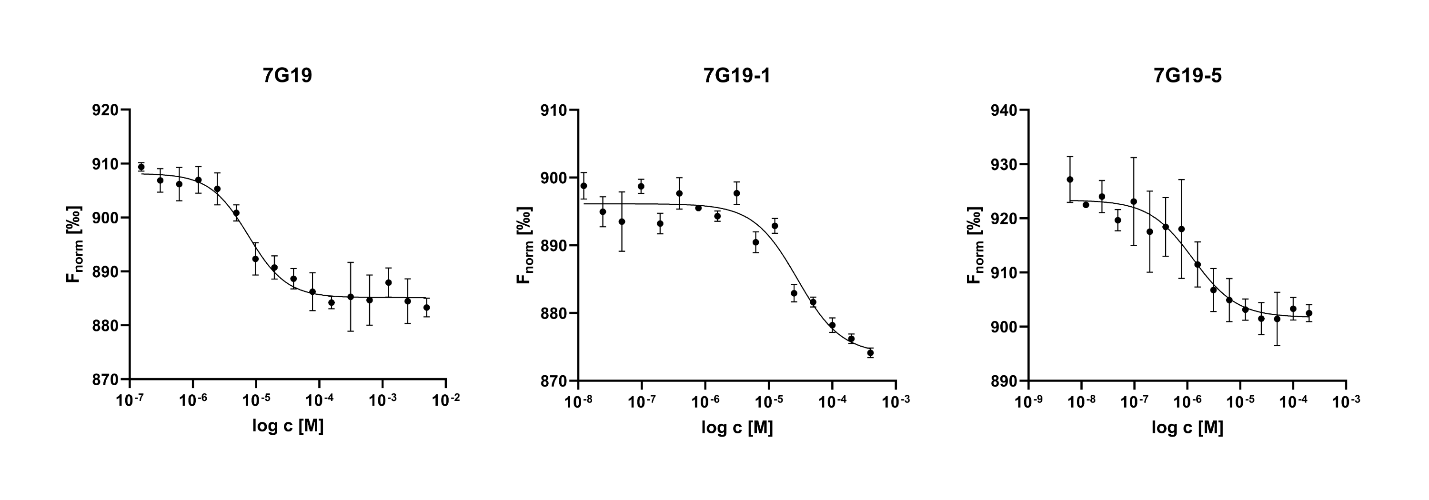

**Figure S9.** F_norm_ vs log c plots for **7G19** derivatives from three independent experiments per compound. Graphs created with GraphPad Prism 10.

#### 2M06 derivatives

**Table S4.** List of 2M06 derivatives with parent compound.

| **well** | **cpd ID** | **MW [g/mol]** | **structure** | ***K*_D_ [µM]** |
| --- | --- | --- | --- | --- |
| 2M06 | Z1443635422 | 394.46 | 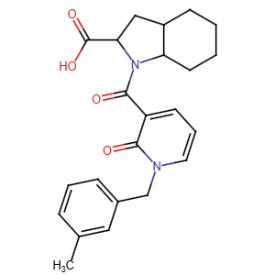 | 26.1 ± 5.1 |
| 2M06-1 | Z841985998 | 339.39 | 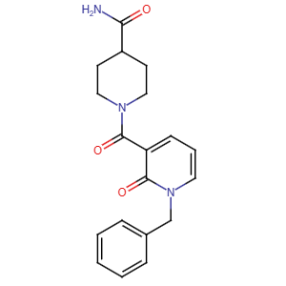 | 124 ± 36 |
| 2M06-2 | Z844993096 | 338.44 | 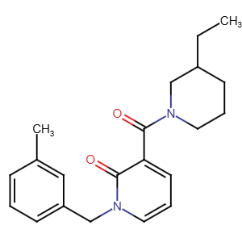 | >500 |
| 2M06-3 | Z841190818 | 310.39 | 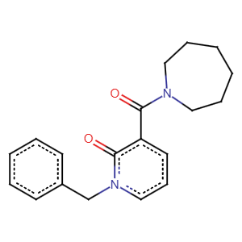 | n. d. |
| 2M06-4 | Z1411892032 | 379.45 | 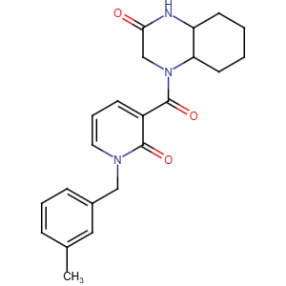 | n. d. |
| 2M06-5 | Z844174818 | 340.42 | 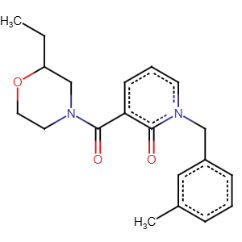 | >500 |
| 2M06-6 | Z1222430106 | 304.34 | 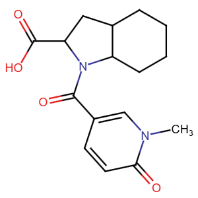 | >300 |
| 2M06-7 | Z970780198 | 352.47 | 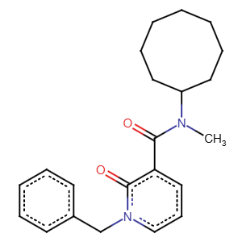 | n. d. |
| 2M06-8 | Z841132532 | 338.44 | 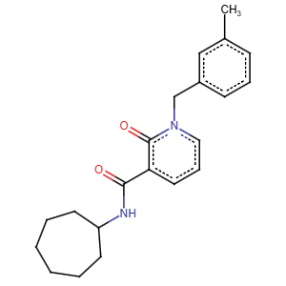 | >300 |
| 2M06-9 | Z2242915781 | 339.43 | 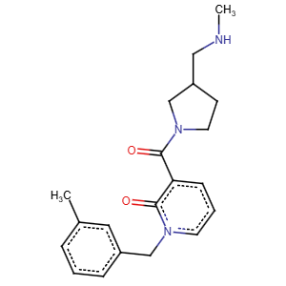 | n. d. |
| 2M06-10 | Z1443621219 | 354.40 | 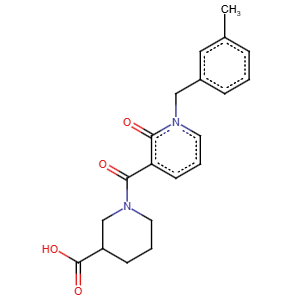 | >500 |

n. d. not determined.

#### 6B10 derivatives

**Table S5.** List of 6B10 derivatives with parent compound.

| **well** | **cpd ID** | **MW [g/mol]** | **structure** | ***K*_D_ [µM]** |
| --- | --- | --- | --- | --- |
| 6B10 | Z1456536982 | 266.77 | 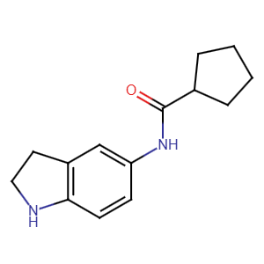 | 37.6 ± 15.8 |
| 6B10-1 | Z1117899740 | 202.25 | 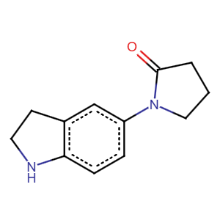 | 19.5 ± 6.8 |
| 6B10-2 | Z1456539098 | 268.78 | 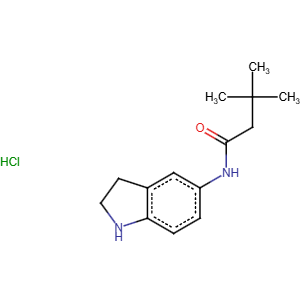 | >300 |
| 6B10-3 | Z3325665525 | 296.79 | 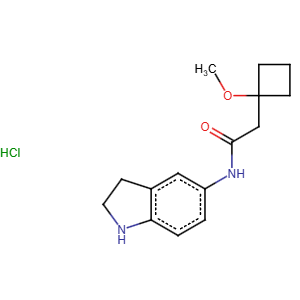 | >300 |
| 6B10-4 | Z73312046 | 272.34 | 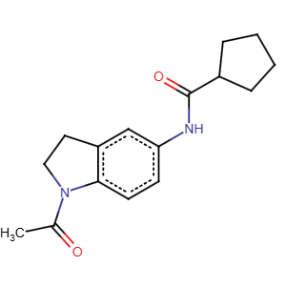 | n. d. |
| 6B10-5 | Z221349054 | 340.37 | 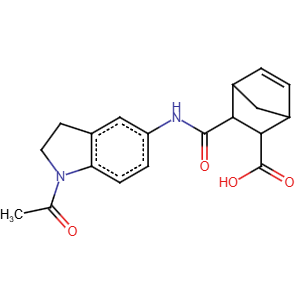 | n. d. |
| 6B10-6 | Z73312179 | 314.42 | 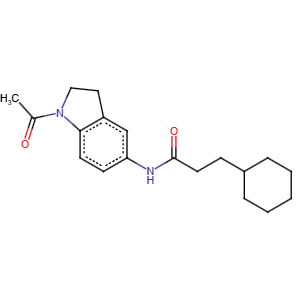 | n. d. |
| 6B10-7 | Z200250388 | 440.56 | 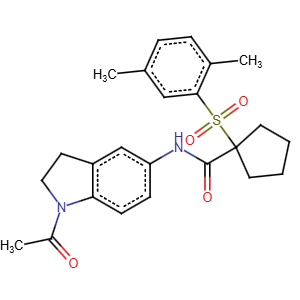 | n. d. |
| 6B10-8 | Z1029824826 | 338.40 | 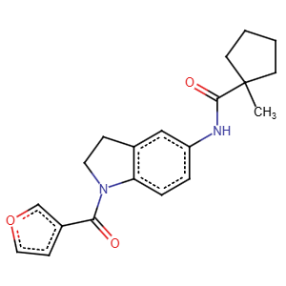 | 136 ± 14 |
| 6B10-9 | Z1095760212 | 348.40 | 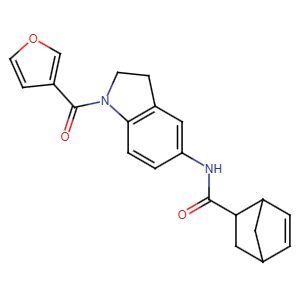 | 68.3 ± 2.8 |
| 6B10-10 | Z1919786242 | 258.32 | 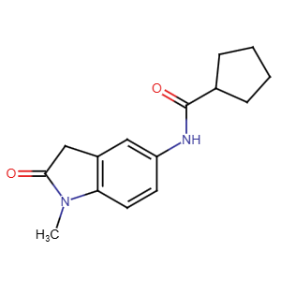 | n. d. |
| 6B10-11 | Z1198959892 | 230.26 | 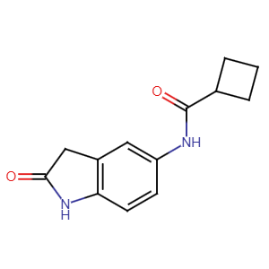 | n. d. |
| 6B10-12 | Z1315014885 | 272.30 | 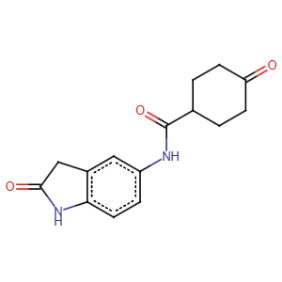 | n. d. |
| 6B10-13 | Z131311418 | 297.35 | 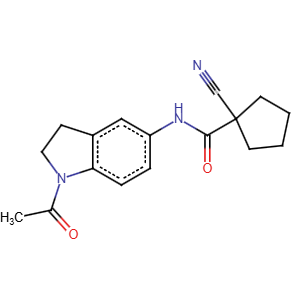 | >300 |
| 6B10-14 | Z1198955791 | 283.33 | 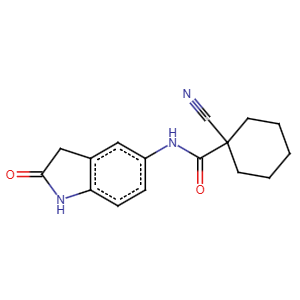 | n. d. |

#### 7G19 derivatives

**Table S6.** List of 7G19 derivatives with parent compound.

| **well** | **cpd ID** | **MW [g/mol]** | **structure** | ***K*_D_ [µM]** |
| --- | --- | --- | --- | --- |
| 7G19 | Z1276736199 | 341.40 | 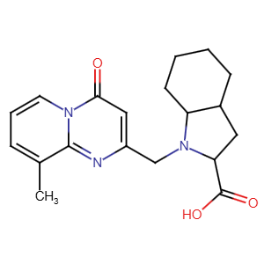 | 3.8 ± 0.3 |
| 7G19-1 | Z1587265793 | 391.51 | 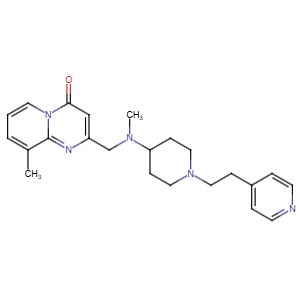 | 70.1 ± 20.0 |
| 7G19-2 | Z840181242 | 313.39 | 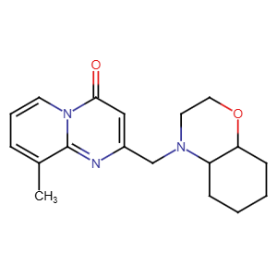 | n. d. |
| 7G19-3 | Z1171370884 | 317.34 | 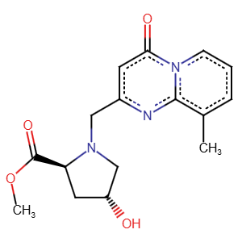 | n. d. |
| 7G19-4 | Z1728138031 | 398.28 | 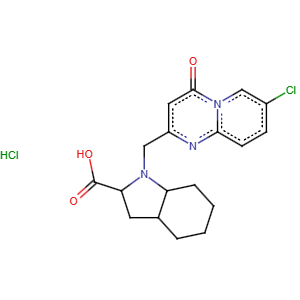 | n. d. |
| 7G19-5 | Z1276735834 | 327.38 | 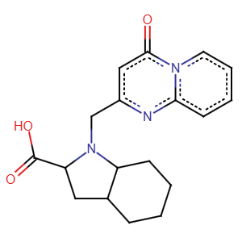 | 4.5 ± 0.9 |

n. d. not determined.

### Thermal shift assay

#### Assay development with positive control T2337

To establish a thermal shift assay for His-TREM2, compound **T2337** was used as a positive control. The assay was conducted with a Prometheus Panta (NanoTemper, Munich, Germany) instrument as described in detail in the materials and methods section. First, assay conditions were optimized without the ligand (**Figure S10**). Addition of 2% DMSO to the assay buffer did not have a significant effect on the protein’s melting temperature (T_m_ = 50.25 ± 0.69 °C in PBST, 49.96 ± 0.42 °C in PBST + 2% DMSO).

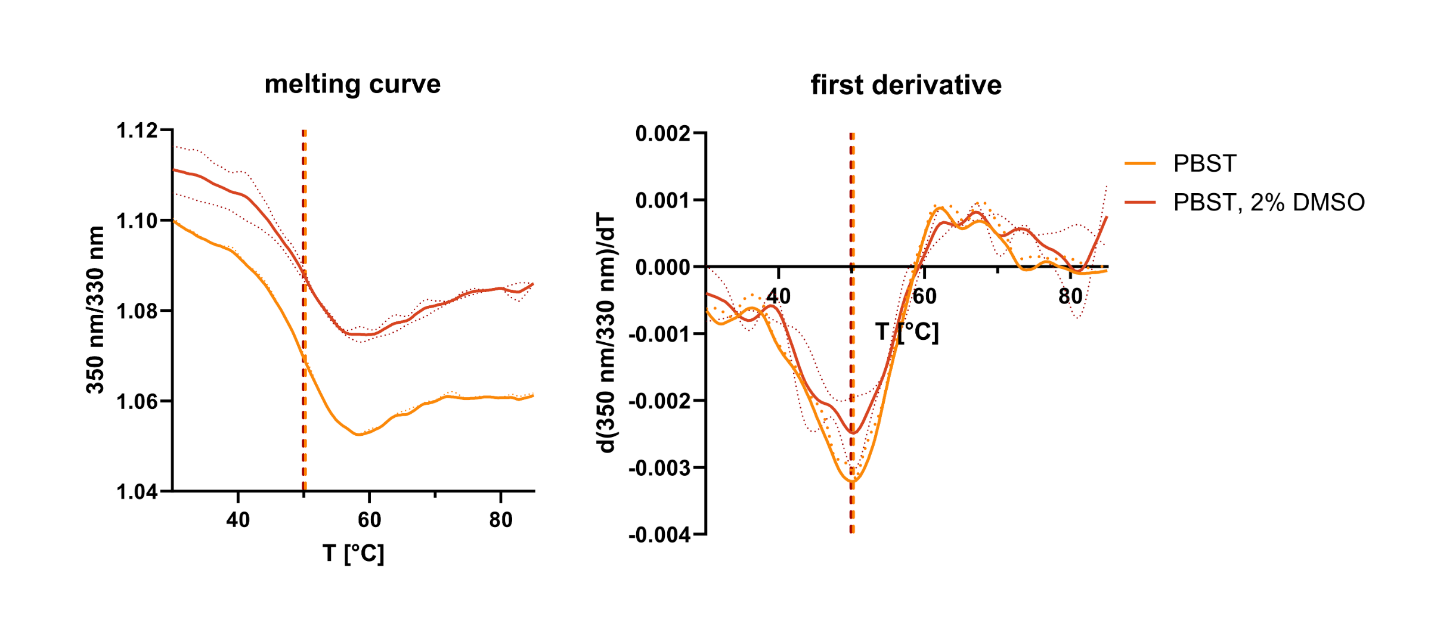

**Figure S10.** Melting curves for His-TREM2 in PBST and PBST with 2% DMSO. Corresponding melting temperatures are indicated as dotted lines. Processed raw data depicted as means with standard deviations from three independent experiments. All graphs created with GraphPad Prism 10.

As a next step, the ligand was used at different concentrations (200–12.5 µM) and preincubated with His-TREM2 for 10 min at rt. A DMSO control was run with the samples. Then, the thermal shift (ΔT_m_) was calculated by the Panta analysis software compared to the control. The melting curves for different **T2337** concentrations are depicted in **Figure S11A** as well as T_m_ vs log c plots (**Figure S11B**), and ΔT_m_ vs log c plots (**Figure S11C**) for better comparability between experiments. Both plots show a dose-dependent thermal shift, resulting in EC_50_ values around 80 µM, which are in the same range as previously determined K_D_ values (MST: 22.4 µM, SPR: 47.8 µM).^1^

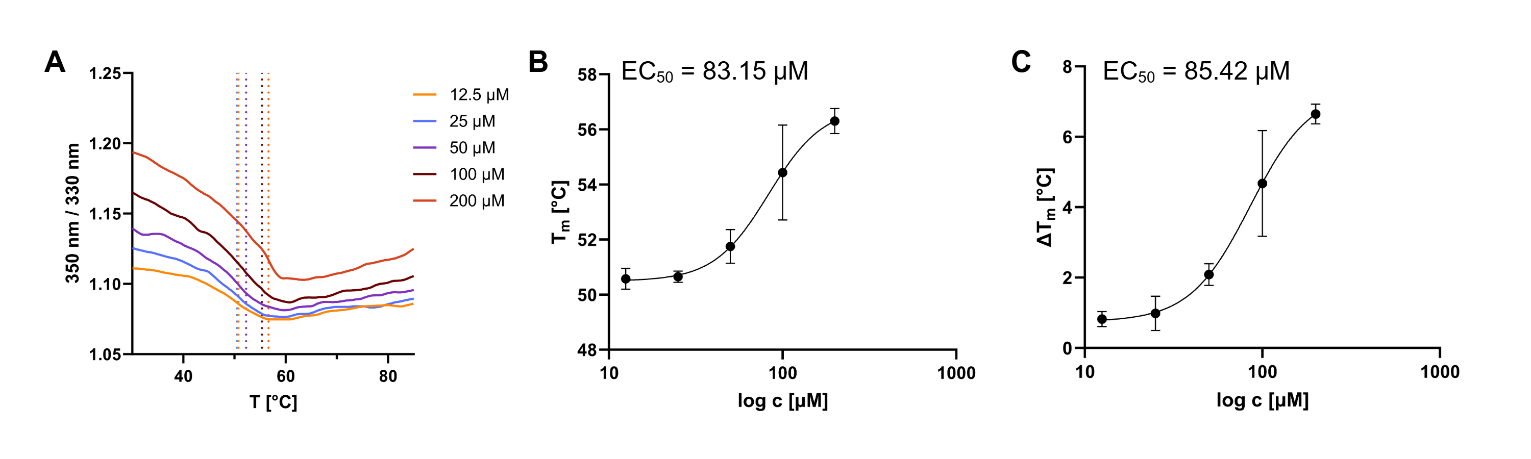

**Figure S11.** Melting profile of His-tagged TREM2 in complex with positive control **T2337**. (**A**) Melting curve of His-TREM2 with **T2337** at different concentrations. Corresponding melting temperatures (T_m_) are indicated as dotted lines. (**B**) Melting temperature plotted against compound concentration to determine EC_50_ value. (**C**) For better comparability between replicates, the difference in T_m_ compared to the DMSO control is plotted against the compound concentration. All graphs were created from raw data with GraphPad Prism 10 and depict the average from three dependent experiments.

#### Single-dose screening

The initially validated hits (**2M06**, **6B10**, **7G19**) as well as selected corresponding derivatives were incubated following the description in the materials and methods section. Compound **T2337** (thermal shift: ΔT_m_ ≈ 5 °C) was run with the samples as a positive control, DMSO in assay buffer as a negative control. The melting curves are displayed as fluorescence ratio vs temperature plots, melting temperatures (T_m_) are indicated as dotted lines (**Figure S12**).

Although nanoDSF is a label-free method that relies on the intrinsic tryptophan fluorescence of the protein, we included a control run with ligands only in assay buffer to exclude the possibility of assay interference (**Figure S12**). Most compounds did not exceed the threshold of 100 fluorescence counts in the discovery scan (exceptions: **2M06-8** and **7G19** for 350 and 330 nm; **7G19-1** and **7G19-5** for 350 nm only), indicating that they would not interfere. However, we selected all capillaries manually to record their melting profiles anyway. Here, some compounds displayed a significantly higher fluorescence ratio than controls (**2M06-8**, **7G19**, **7G19-1**, and **7G19-5**) but their melting graphs did not result in inflection points that could be calculated as T_m_.

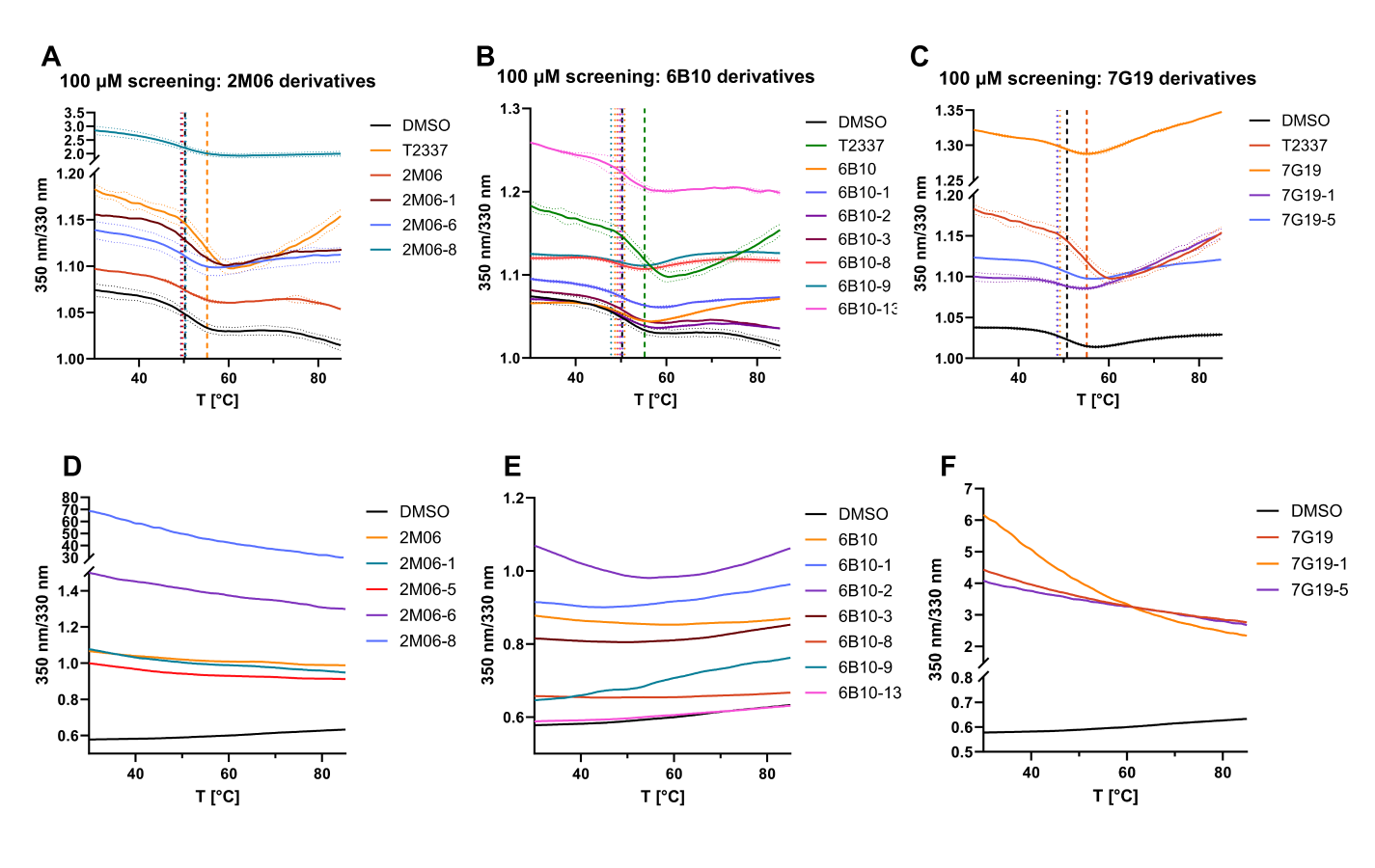

**Figure S12.** Top (**A**–**C**): melting curves for His-TREM2 in complex with hit compounds and selected derivatives in a single-dose screening (100 µM, n = 3). Bottom (**D**–**F**): results from control experiments with ligands only in assay buffer (n = 1) including DMSO control in assay buffer (n = 2).

### Selectivity studies

#### Selectivity against LAG-3

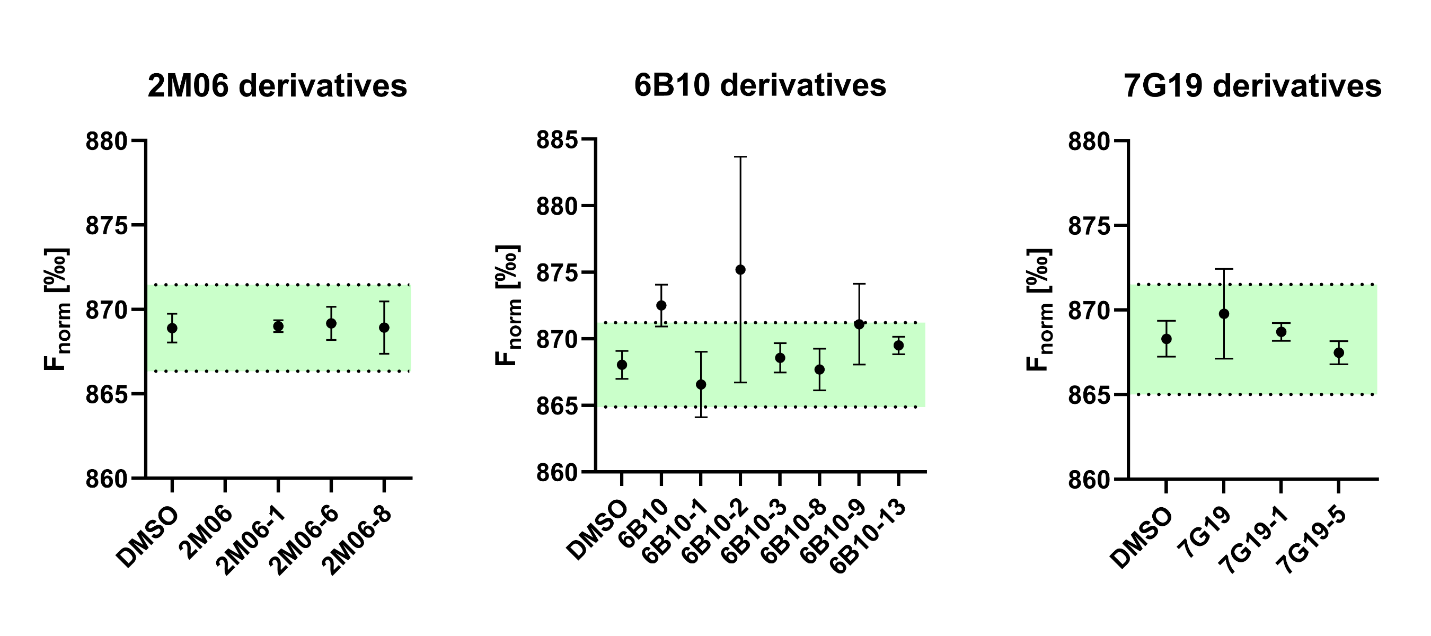

**Figure S13.** Results for LAG-3 selectivity after single-dose screening at 300 µM. All samples and DMSO control are depicted as means with standard deviations. The green area represents three standard deviations from the average control. Graphs created with GraphPad Prism 10.

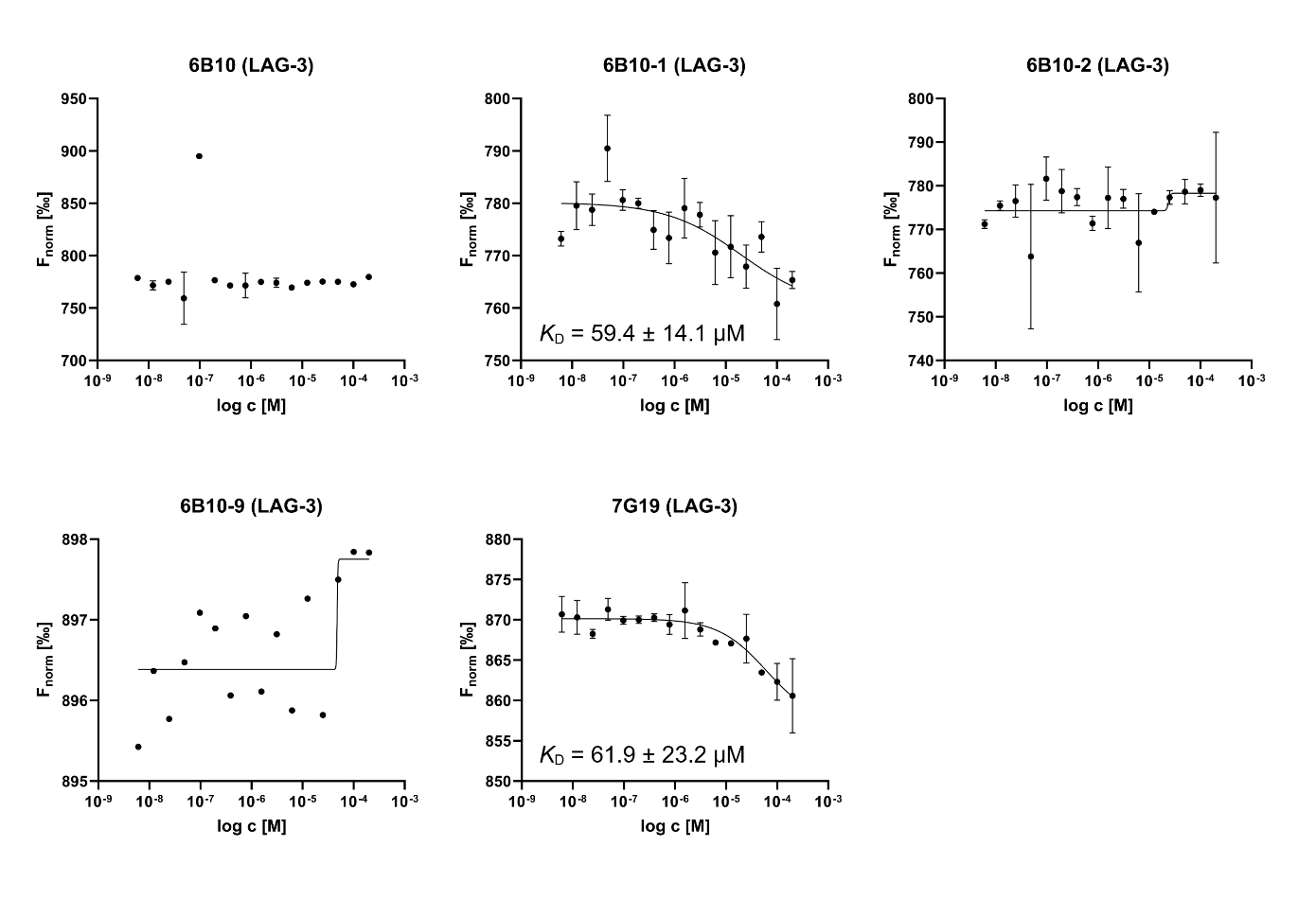

**Figure S14.** Results for LAG-3 selectivity in dose-dependent experiments. Data represented as means with standard deviations. Graphs created with GraphPad Prism 10.

#### Cell-based phagocytosis assay

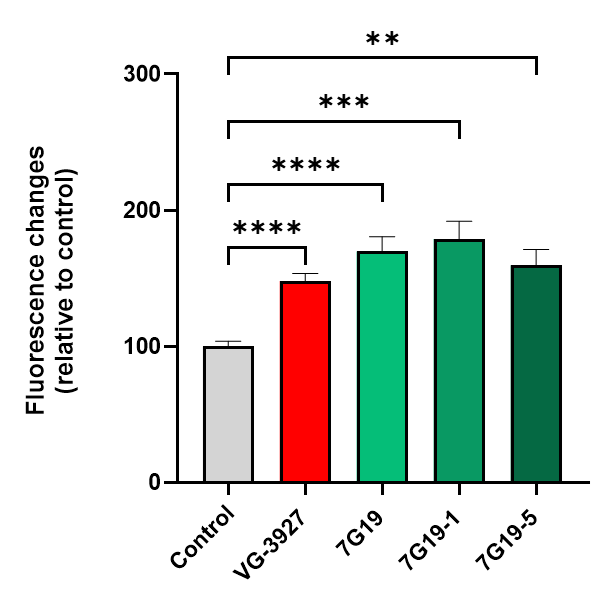

**Figure S15.** Phagocytic activity in HMC3 cells, Histogram representing fluorescence intensity changes, reflecting internalization of pHrodo™ bioparticles. Mean fluorescence values for each condition were normalized to the control group and expressed as relative fluorescence change. HMC3-hTREM2 cells were treated with the indicated compounds (25 μM) for 30 min prior to exposure to pHrodo™ bioparticles for 60 min. **VG-3927** served as a reference, **7G19, 7G19-1** and **7G19-5** display increased phagocytic activity. Data are presented as mean ± SEM from 15 biological replicates. Statistical analyses were performed using Brown–Forsythe and Welch ANOVA tests followed by Dunnett’s T3 multiple comparisons correction.
****p < 0.0001.

SYK and DAP AlphaLisa phosphorylation assay

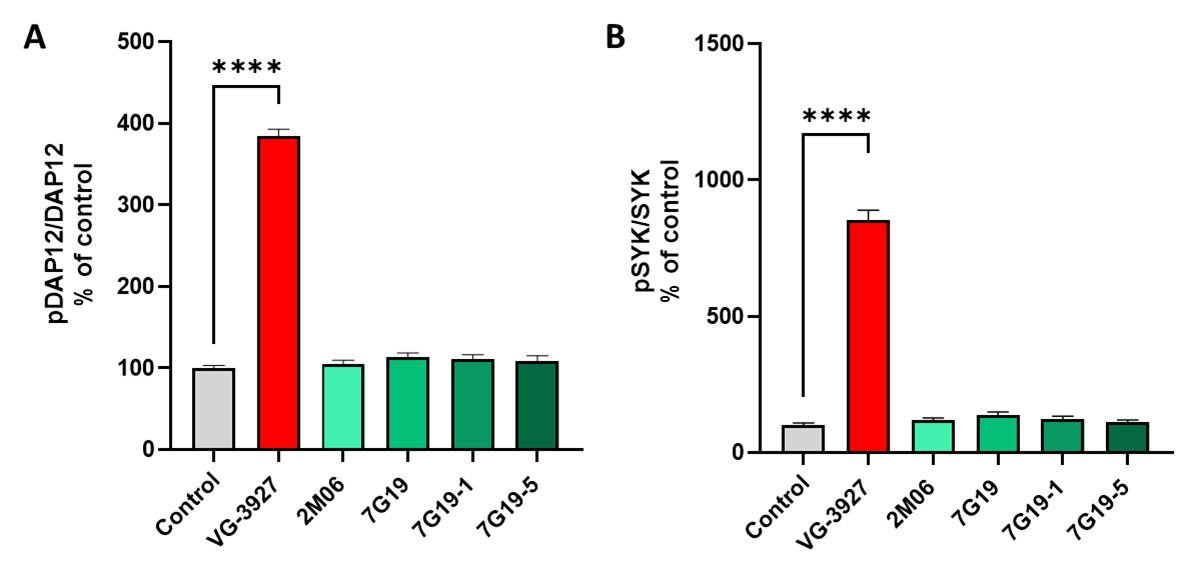

**Figure S16****.** Histogram representing the quantification of phospho-DAP **(A)** and phospho-SYK **(B)** levels, measured using AlphaLisa technique, in untreated condition or after treatment with compound **VG-3927** (serving as reference) or **2M06, 7G19, 7G19-1** or **7G19-5** in TREM2/DAP12 overexpressing HEK cells. Cells were treated with 25 µM of compound for 1 h. Phospho-Syk and phospho-DAP changes in treated conditions are expressed as percentage of control. Data are presented as mean ± SEM from independent biological replicates (n = 4–8). Statistical analyses were conducted using Brown–Forsythe and Welch ANOVA tests, followed by Dunnett’s T3 multiple comparisons correction. Error bars represent means ± SEM. ***p* < 0.005, *****p* < 0.0001.
